# Molecular mechanism of IF1- and IF2-driven translation initiation in bacteria

**DOI:** 10.64898/2026.06.26.734871

**Authors:** Gabriel Soares Guerra, Hassan Zafar, Xueliang Ge, Ritwika S. Basu, Chenhui Huang, Ahmed H. Hassan, Nicolette A. Valdez, Sylva Brabencova, Lucie Slamova, Chandra Sekhar Mandava, Howard Gamper, Ya-Ming Hou, Matthieu G. Gagnon, Suparna Sanyal, Gabriel Demo

## Abstract

Bacterial translation initiation is a highly regulated process essential for accurate start codon selection and the assembly of an elongation-competent ribosome. Two initiation factors, 1 (IF1) and 3 (IF3), contribute to quality control for formation of 30S preinitiation complex (30S PIC), while the GTPase IF2 facilitates stable initiator tRNA binding and promotes subunit association. However, the molecular mechanism of IF1 action and the regulatory role of IF2-mediated GTP hydrolysis and inorganic phosphate (Pi) release remain poorly understood. Using ensemble cryo-EM integrated with fast-kinetics, we delineate the translation initiation pathway involving IF1 and IF2. We show that IF1 transiently associates with the 30S subunit and interferes with the formation of multiple inter-subunit bridges. IF2 promotes subunit association by stabilizing the 30S PIC through interactions mediated by its N-terminal domains. IF1 departure happens after or concomitant with GTP hydrolysis, following which the inter-subunit bridges establish. Then Pi release triggers remodeling of IF2 followed by its departure from the 70S initiation complex. These findings reveal how the coordinated interplay of IF1 and IF2 with the ribosome ensures translational fidelity and plays crucial role for formation of elongation-competent 70S.

## Introduction

Translation initiation is a critical control point in protein synthesis, ensuring that the ribosome assembles at the canonical start codon and in the proper conformation to enter the elongation phase of translation^1–3^. In bacteria, this multi-step process is coordinated by three initiation factors – IF1, IF2, and IF3 – which guide the selection and positioning of the initiator tRNA (fMet-tRNA^fMet^), promote accurate start codon–anticodon pairing, and drive the transition from a 30S preinitiation complex (30S PIC) to a 70S elongation-competent (70S EC) ribosome^4^.

Initiation factors IF1 and IF3 play essential and complementary roles in ensuring the fidelity and directionality of bacterial translation initiation^5–8^. IF1 binds to the aminoacyl (A) site of the 30S small ribosomal subunit, where it prevents premature entry of elongator tRNAs by shielding the mRNA codon and ensures binding of fMet-tRNA^fMet^ in the peptidyl (P) site^9,10^. IF1 also interacts with the decoding centre (DC) and has been shown to inhibit formation of inter-subunit bridge B2a, thereby affecting 50S large ribosomal subunit joining^5,9–11^. In parallel, IF3 binds near the exit (E) site of the 30S subunit and is critical for accurate fMet-tRNA^fMet^ selection^5,12^. It prevents premature subunit joining by interfering with the formation of key inter-subunit bridges and facilitates proofreading of 30S PICs^9,10^, although structural studies of later initiation stages in its absence have demonstrated stable 70S complex formation^11^. The dynamic interaction of IF3 with the 30S subunit^10^, in cooperation with IF1, enhances the correct positioning of fMet-tRNA^fMet^ within the 30S PIC^5^. Together, IF1 and IF3 flank fMet-tRNA^fMet^, promoting its correct base pairing with the AUG start codon in the mRNA positioned at the P site accompanied by the closure of the 30S subunit head domain, which in turn locks the fMet-tRNA^fMet^ in place within the 30S PIC^9,10^. Beyond this structural role, IF1 and IF3 serve as quality control checkpoints, discriminating against improperly assembled 30S PICs and, when necessary, blocking subunit joining^12,13^. IF2, the largest and most complex of the bacterial initiation factors, takes a central role in initiation^11,14–16^ as the C2-domain of IF2 stabilizes fMet-tRNA^fMet^ by specifically recognizing the formyl-methionine (fMet) moiety at the 3′-CCA end of the fMet-tRNA^fMet 10,11^. This interaction not only reinforces the correct positioning of the fMet-tRNA^fMet^ in the P site but also promotes the release of IF3, setting the stage for 50S subunit joining^10,11^. However, beyond occluding the A site, the molecular mechanism of IF1-mediated discrimination against incorrectly formed 30S PICs remains unclear. Early time-resolved cryo-EM and kinetic studies revealed IF1 blocking the inter-subunit bridge B2a, suggesting IF1 dissociates rapidly after subunit joining but before dissociation of IF2^8,11^. The role of IF1 in coordination with IF2 as a fidelity-checkpoint has been proposed as critical for translation initiation quality control^12,13^, but the mechanism has remained unknown.

IF2 is a GTPase that accelerates subunit joining^17^ and mediates fMet-tRNA^fMet^ accommodation into the P site on the 50S subunit, thereby promoting the formation of 70S IC. In *E. coli*, IF2 exists in multiple isoforms (α, β, and γ) arising from alternative translation initiation sites, with the γ-isoform also arising from proteolytic cleavage of the α-isoform (IF2α)^18,19^. Under normal growth conditions, IF2α is predominantly active and enhances subunit joining and initiation efficiency^20^, whereas the shorter variants become active under stress conditions such as cold shock^21,22^, underscoring distinct roles for each isoform. While all IF2 isoforms share the conserved GTPase domain (G-domain) and tRNA-binding core, β/γ-isoforms lack the extensive N-terminal extension found in the α-isoform^23,24^. The N2-domain has been implicated in stabilizing IF2 association with the 30S subunit via contacts with helix 16 (h16) of 16S rRNA^20,25,26^ independently of both GTP/GDP binding and the presence of IF1^27^. In contrast, the γ-isoform in *B. stearothermophilus* requires both GTP and IF1 for stable 30S association^27^. Strikingly, recent work shows that IF2α interacts directly with the transcription factor NusA, suggesting that its N-terminal extension may play additional regulatory roles^28^, although its structural contribution to ribosome binding remains unclear.

The overarching function of IF2 is to facilitate the full accommodation of fMet-tRNA^fMet^ into the P/P state, a key step of the transition from 70S IC to 70S EC^11,16,26,29,30^. This transition is triggered by GTP hydrolysis at IF2’s G-domain, catalysed by His448 within the switch II loop (sw-2) and coordinated by residues from switch I region (sw-1), the sarcin-ricin loop (SRL) in the 50S subunit, and a magnesium ion that coordinates with the nucleotide^11,12,14,29,31^. A recent structural study in *Pseudomonas aeruginosa* revealed a dramatic rearrangement of GDP-bound IF2α into a compact conformation, a structural intermediate resulting from the remodelling of sw-2 which progressively relieves constraints on the C1-domain of IF2α, effectively pulling the C2-domain away from the 3′-CCA end of fMet-tRNA^fMet^ to facilitate its full accommodation into the P/P state^26^. Yet, the interplay between GTP hydrolysis, Pi release, and IF2 conformational dynamics during translation initiation remains elusive.

Here, we combine ensemble cryo-EM with fast-kinetics to capture structurally distinct intermediates along the *E. coli* translation initiation pathway, spanning the transition from the 30S PIC to the 70S EC through GTP hydrolysis-driven maturation of the 70S IC. Upon formation of the 70S IC, IF1 interferes with the formation of four inter-subunit bridges, including B2a, while IF2α compensates for reduced inter-subunit contacts in early 70S ICs through interactions of its core domains with the 50S subunit. Subsequent GTP hydrolysis by IF2α and inorganic phosphate (Pi) release induce large conformational changes in IF2α that ultimately lead to its dissociation from the ribosome. Our work thus provides novel structural and mechanistic insights into key events of bacterial translation initiation, revealing the interplay between IF1 and IF2α in ensuring quality control and outlining a stepwise, GTP hydrolysis-driven pathway for translation initiation.

## Results

### IF2α promotes rapid subunit association

To characterize the role of the N-terminal domains (NTDs) of IF2α, we prepared NTD-truncated mutant constructs, corresponding to deletions D1 (Δ1-159, also known as the β isoform) and D2 (Δ1-289) (Fig. 1a). All constructs contained the N2-domain but varied in the N1-domain. We first examined the activity of the IF2 variants through [³H]-GTP binding assays. All variants exhibited a similar linear increase in GTP binding across the tested concentration range, indicating that they are active and fully bound to GTP in our functional assays (Supplementary Fig. 1a, b). We have also checked whether the NTD truncation of IF2 affects binding of the initiator tRNA. Using fluorescent BOP-Met-tRNA^fMet^ we show that all three IF2 variants support initiator tRNA binding to the ribosome in a similar manner (Supplementary Fig. 1c).

**Figure 1.**
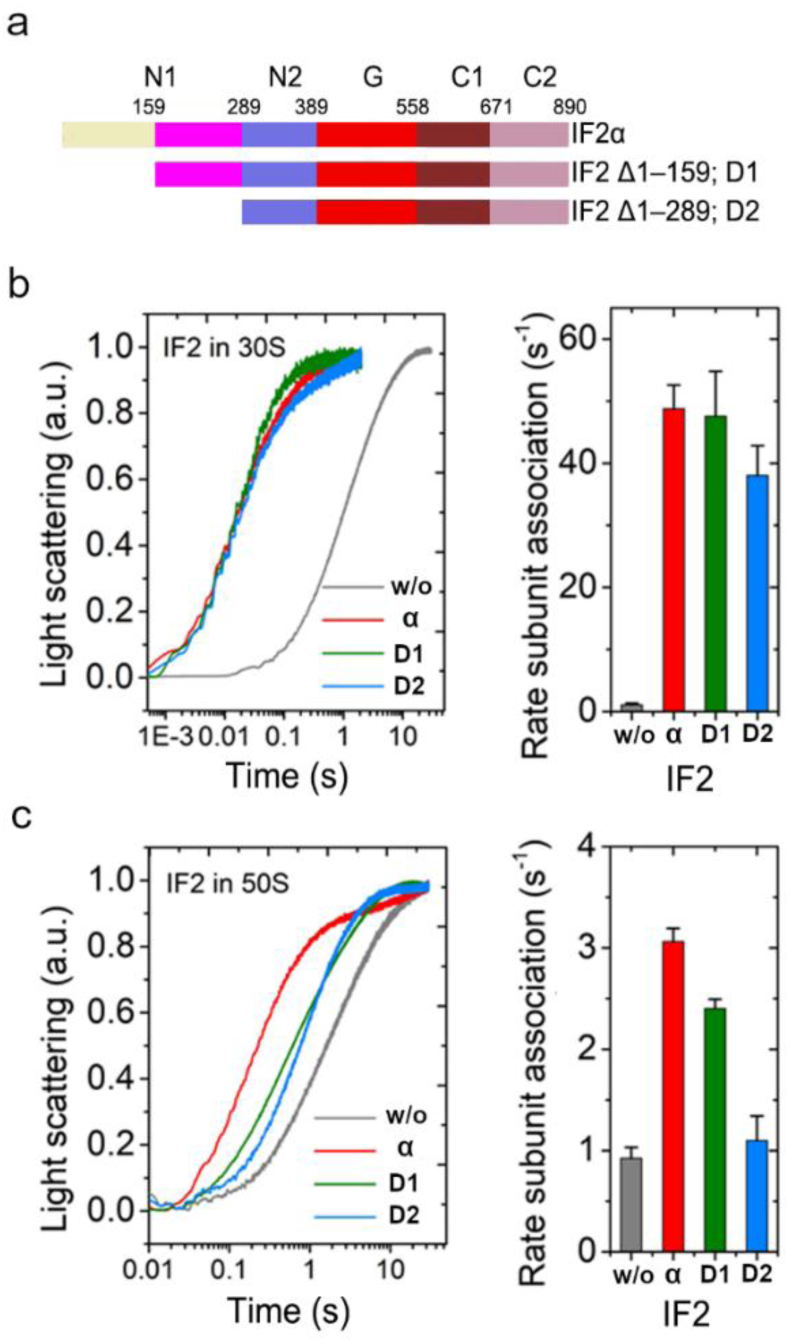
Comparison of IF2α, D1 and D2 variants in subunit association. (**a**) Schematic representation of three IF2 variants specifying respective N-terminal deletions. Kinetics of subunit association followed by Rayleigh light scattering in stopped-flow with IF2 variants, either pre-equilibrated with 30S PIC (**b**) or supplied with 50S subunit (**c**). The right panels demonstrate the rates of subunit association for each IF2 construct. Data are presented as mean ± s.d. from three independent experiments.

Next, IF2α, D1, and D2 were subjected to subunit association assays, in which 50S subunits were rapidly mixed with the 30S PIC using a stopped-flow instrument, and the increase in Rayleigh light scattering over time was monitored. IF2 was added either to the 30S PIC mix or with the 50S subunits. While all IF2 variants facilitated rapid subunit association, no significant differences among the three were observed when IF2 was pre-incubated with 30S. In comparison to a rate of 0.9 ± 0.1 s⁻¹ without IF2, IF2α and D1 increased the subunit association rate to approximately 50 s⁻¹, while D2, with its larger truncation, exhibited a slightly slower rate of 40 s⁻¹ (Fig. 2b). Since D1 and D2 variants lack different length of N1-domain, it can be concluded that the N1-domain is not essential for efficient subunit association, when the factor is stably bound to 30S PIC.

**Figure 2.**
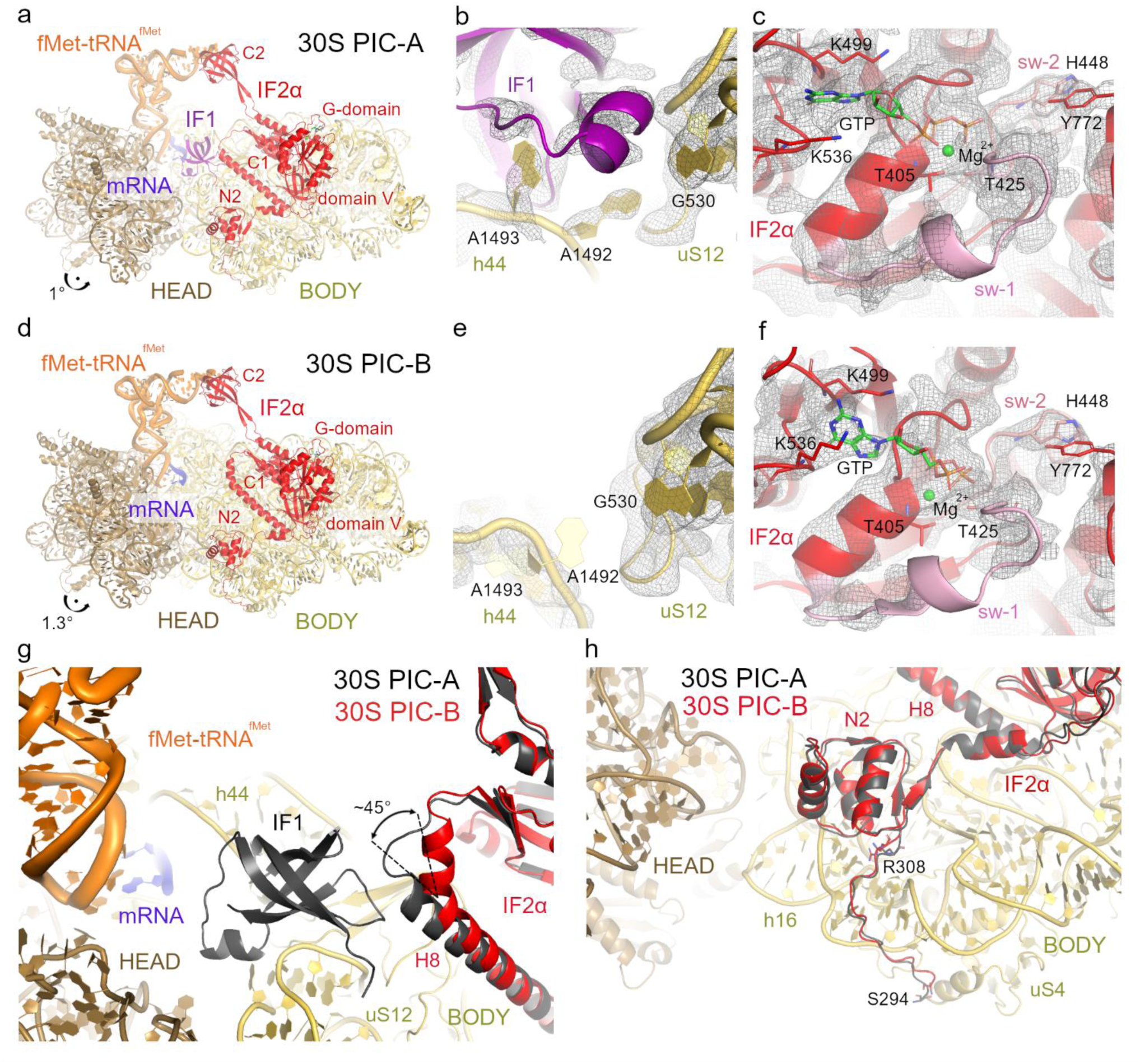
Cryo-EM structures of the 30S PICs. (**a**) Overall structural view of the 30S PIC-A complex. (**b**) Close-up view of the decoding centre (DC) in 30S PIC-A, highlighting the stabilization of nucleotides A1492 and A1493 by IF1. The original cryo-EM density map is shown as a grey mesh at 3.0 σ. (**c**) Detailed view of the G-domain of IF2α in 30S PIC-A. The switch I (sw-1) and switch II (sw-2) regions are shown in light pink and salmon, respectively. The corresponding cryo-EM density is displayed in grey mesh at 1.8 σ. (**d**) Overall structural view of the 30S PIC-B complex. (**e**) Close-up view of the DC in 30S PIC-B. The original cryo-EM density map (3.0 σ, grey mesh) indicates high flexibility of the DC nucleotides in h44 of the 16S rRNA. (**f**) Detailed view of the G-domain of IF2α in 30S PIC-B. The sw-1 and sw-2 regions are coloured as in panel (c). The cryo-EM density is displayed in grey mesh at 1.6 σ. (**g**) Structural alignment of 30S PIC-A (dark grey) and 30S PIC-B (coloured as in panel d), based on the 16S rRNA of the 30S body from 30S PIC-B. IF1 is stabilized by the C-terminal portion of IF2α helix 8 (H8), which adopts a ∼45° kink in its absence. (**h**) Structural alignment showing docking of the IF2α N2-domain onto 16S rRNA helix 16 (h16) near ribosomal protein uS4. The preceding loop (Ser294–Arg308) is highlighted. In panels (a) and (d) IF2α is shown in red, IF1 in purple, 30S head in light brown, 30S body in yellow, fMet-tRNA^fMet^ in orange, mRNA in dark blue, GTP (sticks) and Mg^2+^ (sphere) in green. Arrows indicate the 30S head swivel in panels (a) and (d).

The subunit association reactions, however, slowed considerably when IF2 was added with the 50S subunits, indicating that the reaction was limited by IF2 binding. Notably, now the IF2α, D1, and D2 variants exhibited different rates: the reaction was fastest with IF2α (3.06 ± 0.13 s⁻¹), followed by D1 (2.4 ± 0.09 s⁻¹), and the slowest with D2 (1.1 ± 0.24 s⁻¹), suggesting that IF2α is most efficient in binding to the 30S PIC (Fig. 2c) than D1 and D2, both lacking parts of the N1-domain. A similar trend of IF2α > D1 > D2 was observed when the experiment was repeated with IF3 added to the 30S PIC, although all rates decreased considerably (Supplementary Fig. 2). This finding is supported by earlier studies showing that deletion of just the first 69 residues from IF2α causes a marked reduction in 50S association, impairing proper 70S IC formation^20^. Consistently, single-molecule FRET analyses have shown that IF2α promotes a productive conformation of the 30S PIC, favouring rapid 50S recruitment and efficient 70S IC assembly^20,30^.

These results demonstrate that the N1-domain of IF2 does not directly affect the mechanical process of 50S subunit docking. Instead, truncations of the N1-domain in D1 and D2 slow their recruitment to the 30S PIC, leading to kinetic penalties during subunit association. Collectively, these findings identify the IF2 NTDs as a critical determinant for the efficient and stable association of IF2 with the 30S PIC, thereby influencing the overall rate of translation initiation.

To investigate the early stages of bacterial translation initiation, we designed an ensemble cryo-EM approach that exploits the kinetic properties of IF2α-dependent subunit joining (Fig. 1b, c). Accordingly, IF2α was supplied only with the 50S subunit together with GTP, slowing the transition toward 70S IC formation. To enable accumulation of 70S ICs, IF3 was omitted because of its strong anti-association activity^5^. Consequently, we obtained an ensemble enriched in both unjoined 30S PICs and 70S ICs, enabling structural analysis of the coordinated roles of IF1 and IF2 during subunit joining and 70S IC formation, while excluding IF3-dependent quality-control mechanisms during translation initiation.

### IF1 binding to 30S PICs and anchoring role of IF2α N2-domain

We resolved two distinct 30S PIC structures, termed 30S PIC-A and 30S PIC-B, both assembled with IF2α and fMet-tRNA^fMet^ in the P/I configuration (Fig. 2; Supplementary Table 1, Supplementary Figs. 3 and 4). In both complexes, the AUG start codon of the mRNA is base-paired with the anticodon of fMet-tRNA^fMet^ (Supplementary Fig. 5a, b), indicating canonical codon–anticodon recognition in the P site and the adoption of the closed conformation of the 30S subunit previously associated with full accommodation of the initiator tRNA in the P/I state^10,11^. IF2α is clearly resolved in both structures, with its C2-domain contacting the 3′-CCA end of fMet-tRNA^fMet^ (Fig. 2a, d). Although this interface appears flexible and locally less well defined likely due to the absence of stabilizing effect of the 50S subunit, focused 3D classification revealed conformational variability in this region (Supplementary Fig. 6). By contrast, the G-domain of IF2α is well resolved, allowing confident placement of GTP (Fig. 2c, f). Additionally, the sw-1 region adopts an ordered conformation in both complexes, consistent with a pre-hydrolysis state^11,26^, as previously observed in similar GTP-bound forms of other translational GTPases, including elongation factors EF-Tu and EF-G^32,33^ (Fig. 2c, f). The catalytic residue His448, located within the sw-2 region of IF2α, is oriented away from the γ-phosphate of GTP, indicating that it is not positioned for catalysis. This misalignment is due to the absence of the 50S subunit, and specifically the sarcin-ricin loop (SRL), which plays a critical role in stabilizing the GTPase active site and properly positioning His448 for hydrolysis (Fig. 2c, f).

The distinctive difference between the two 30S PICs is the presence (30S PIC-A; Fig. 2a) or absence of IF1 (30S PIC-B; Fig. 2d). In 30S PIC-A, IF1 occupies the A site, anchored between helix 44 (h44) of the 16S rRNA and ribosomal protein uS12, and directly contacts the decoding nucleotides A1492 and A1493, which are both observed in flipped-out conformations (Fig. 2b). In 30S PIC-B, in the absence of IF1, A1493 adopts a more flipped-in position, while A1492 remains flipped out towards the A site. However, the local density in this region is weak, indicating high flexibility of the decoding centre (DC) upon IF1 dissociation (Fig. 2e). This observation suggests that IF1 stabilizes h44, and its release may allow local rearrangements of the DC nucleotides. Comparison of the two structures reveals a subtle change in the 30S head region (Fig. 2a, d), likely reflecting local rearrangements rather than a global head swivel, and suggesting that IF1 binding may be sensitive to conformational variability in this region, as reported previously^34^. IF1 is shallowly anchored between h44 and uS12, and its binding likely fluctuates with the intrinsic flexibility of the 30S subunit before 50S joining. In both structures, helix H8 of IF2α interacts with ribosomal protein uS12, which is also a contact site for IF1 (Fig. 2g). Notably, in 30S PIC-B where IF1 is absent, the C-terminal segment of H8 adopts a distinct conformation, forming a ∼45° kink and extending its α-helix by three residues compared to its structure in 30S PIC-A (Fig. 2g). This rearrangement likely represents a structural adaptation of H8 in response to the presence or absence of IF1 in the A site, enabling H8 of IF2α to reposition itself accordingly on the 30S subunit. Together, these structures provide a rationale for earlier biochemical findings^5,29^ showing that *in vitro*, IF2 alone is sufficient to promote 50S subunit joining and formation of a functional 70S IC, while IF1 is dispensable for subunit joining and instead acts to stabilize the DC and modulate the efficiency and fidelity of initiation, consistent with both 30S PIC-A and 30S PIC-B serving as productive substrates for 70S IC formation.

Moreover, our 30S PIC structures resolved IF2 from residue Ser294 onward, revealing the N2-domain in substantial detail. In both complexes, the N2-domain is stably docked into the minor groove of 16S rRNA helix 16 (h16) (Fig. 2h), consistent with previous observations in *P. aeruginosa*^26^ and recent 30S translation initiation complex coupled to paused RNA polymerase^35^. This supports the previous observations that the N-terminal region contributes to stabilizing IF2 interaction with the 30S subunit^20^. Residues upstream of Ser294 were not resolved, consistent with a flexible α-specific N-terminal extension. These observations suggest that stabilization of IF2 on the 30S subunit is primarily mediated by the N2 domain (Fig. 1b, c), while the unresolved α-specific N-terminal region may provide architectural support to the N2 region thereby facilitating this interaction. This is supported by our biochemical data that when pre-mixed with 30S PIC, all IF2 variants containing the N2 domain showed similar rate of subunit association (Fig. 1b).

### IF1 and IF2α in the extended conformation safeguard the formation of the 70S IC

Rapid-kinetic and FRET studies show that blocking IF2-mediated GTP hydrolysis prevents IF1 release from the 70S IC, and that IF1 dissociation coincides with GTP hydrolysis rather than subunit joining^8^. To evaluate the role of IF1 in 70S IC formation and to capture this transient initiation intermediate, we reconstituted the 70S IC from purified subunits in the presence of the non-hydrolysable GTP analogue 5’-guanosyl-methylene-triphosphate (GDPCP) instead of GTP, which can shift the equilibrium toward a stable IF1-bound 70S IC^8^ (Supplementary Table 1; Supplementary Figs. 4 and 7). This yielded a distinct structure, designated as 70S IC* (4.1° body rotation and 1.0° head swivel), capturing an early post-subunit association intermediate in which IF1 is retained (Fig. 3a, b). In the 70S IC*, IF1 occupies its canonical binding site between h44 of the 16S rRNA and uS12, where it stabilizes the decoding nucleotides A1492 and A1493 (Fig. 3b), and forms weak contacts with the C-terminal region of IF2α helix H8, as also observed in the 30S PIC-A structure (Figs. 2g and 3a, b). The IF2α adopts a fully extended conformation^11,26^ (Fig. 3a), similar to that observed in the 30S PICs. The distal C2-domain from IF2α securely engages fMet-tRNA^fMet^ in the P/PI state, while codon–anticodon pairing with the AUG codon is maintained in the P site (Supplementary Figs. 5c and 8a). The presence of the 50S subunit enhances the stability of the interface between the C2-domain of IF2α and the 3′-CCA end of the initiator tRNA (Extended Data Fig. 8a). This stabilization is reflected in the high local resolution of the region and enables clear visualization of critical molecular contacts, where Phe804 and Phe848 flank and stabilize the fMet moiety, while Arg846 envelops the A76 nucleotide of the tRNA, securing the 3′-end within the P/PI state (Extended Data Fig. 8a). Moreover, the G-domain is well resolved, revealing bound GDPCP and an ordered sw-1 region consistent with a locked pre-hydrolysis state (Fig. 3c).

**Figure 3.**
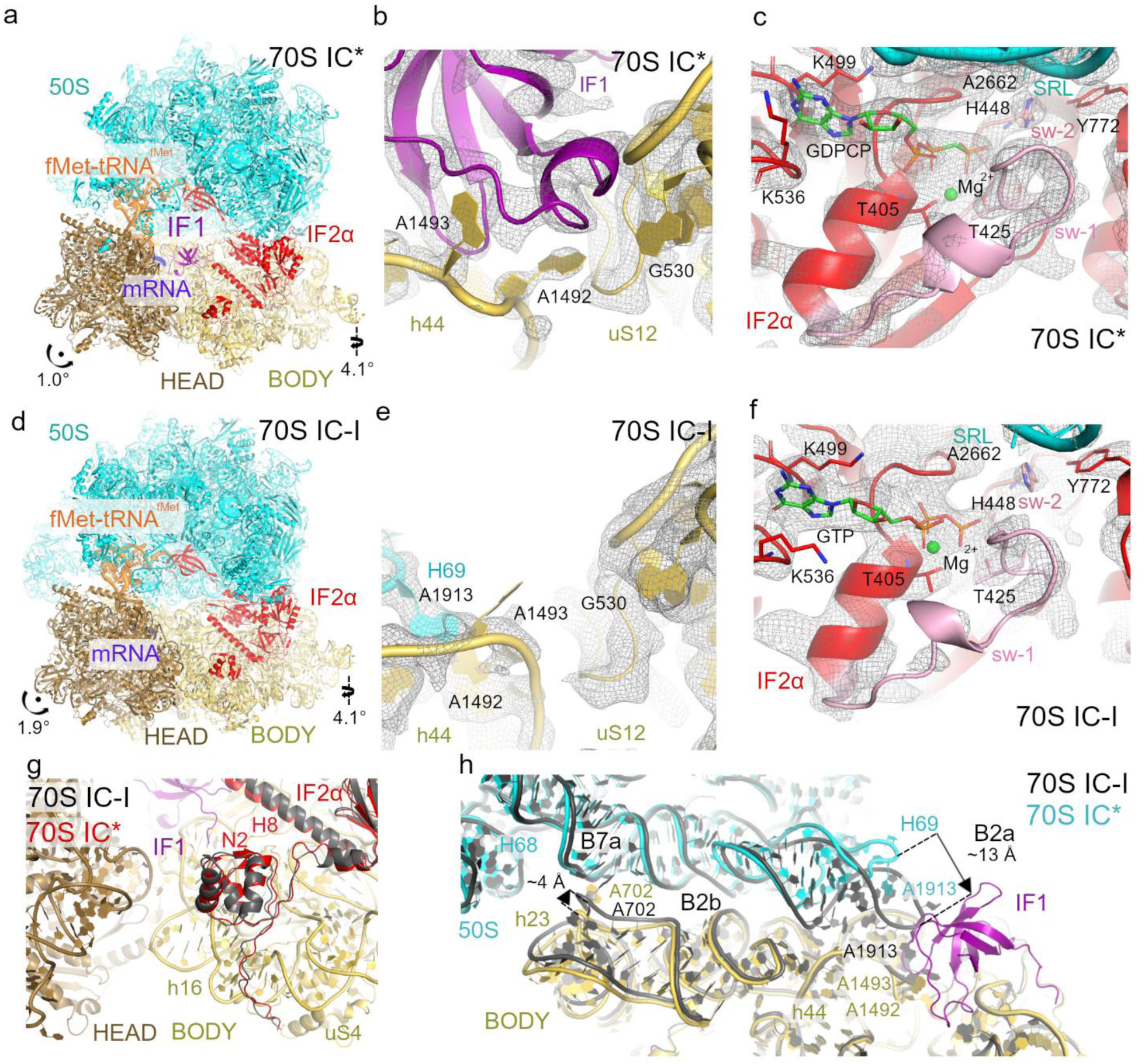
Cryo-EM structures of 70S IC* and 70S IC-I. (**a**) Overall view of 70S IC*, coloured as in Fig. 2, with the 50S subunit shown in cyan. Arrows indicate the 30S head swivel and body rotation. (**b**) Close-up view of the DC in 70S IC*, highlighting the positions of DC nucleotides A1492 and A1493. The original cryo-EM density map is shown as a grey mesh at 2.5 σ. (**c**) G-domain of IF2α in 70S IC*. The purine base of GDPCP stacks against Lys499 and Lys536. Switch I (sw-1, light pink) and switch II (sw-2, salmon) regions are shown. The cryo-EM density is shown as a grey mesh at 2.2 σ (IF2α) and 4.0 σ (SRL). (**d**) Overall view of the 70S IC-I. Arrows indicate the 30S head swivel and body rotation. (**e**) Close-up view of the DC in the 70S IC-I, highlighting the positions of the DC nucleotides A1492 and A1493. The original cryo-EM density is shown as a grey mesh at 2.2 σ. (**f**) G-domain of IF2α in 70S IC-I in presence of GTP. Sw-1 and sw-2 regions are shown in the same colouring as in (c). The cryo-EM density map is shown as a grey mesh at 2.5 σ. (**g**) Structural alignment showing N2-domain of IF2α docked onto h16 of the 16S rRNA in both 70S IC* and 70S IC-I. (**h**) Structural alignment reveals the position of IF1 in the 70S IC*, showing that inter-subunit bridges B2a and B7a remain incompletely formed. Structural alignments of 70S IC-I (dark grey) and 70S IC* are based on the 16S rRNA of the 30S body.

To compare the 70S IC* structure with an on-pathway 70S initiation complex, we performed extensive 3D-focused classification under GTP conditions (Supplementary Table 1; Supplementary Figs. 4 and 9), which revealed three distinct 70S structural classes. The first structure, designated as 70S IC-I, represents a closely related state in which IF1 is absent, while IF2α remains in an extended conformation^11,26^. In 70S IC-I, the 30S subunit exhibits a body rotation of 4.1° relative to the 50S subunit, along with a 1.9° head swivel (Fig. 3d). Notably, comparison with previous structural studies of 70S ICs^11,26^ reveals a similar rotational state of 30S body (Supplementary Fig. 10a, c), supporting the designation of 70S IC-I as an on-pathway intermediate in translation initiation. In this structure, the absence of IF1 correlates with reduced local resolution in h44 of the 16S rRNA, indicative of increased flexibility in the DC (Fig. 3e). Within this region, the decoding nucleotide A1492 is likely flipped-in, stacking against A1913 of helix 69 (H69) in the 23S rRNA, while A1493 remains flipped out towards the A site, bringing the DC into a conformation closer to that observed in elongation-ready 70S ribosomes^11^. The C2-domain engages the 3′-CCA end of fMet-tRNA^fMet^ similarly as in 70S IC* (Supplementary Fig. 8a, b), while canonical base pairing between the tRNA anticodon and the mRNA AUG codon in the P site is preserved (Supplementary Fig. 5d). Although the C2-domain exhibits slight positional variability compared to previous studies^11,26^ (Supplementary Fig. 10a-c), its conformation is overall more stabilized in 70S IC* and 70S IC-I structures than in the 30S PIC structures. At the G-domain centre of 70S IC-I, the cryo-EM density is most consistent with a GTP-bound pre-hydrolysis model (Fig. 3f). However, the geometry of the G-domain active site also closely resembles GDP–Pi states observed in translational GTPases^32,33^ and Ras-family GTPases captured with GDP and inorganic phosphate (Pi)^36,37^ (Supplementary Fig. 11). Positioning of the β-(GDP) and γ-phosphates (Pi) indicates that their relative spacing is comparable to that observed in GDP–Pi bound GTPases, while preserving the coordination sphere of the Mg^2+^ ion required for catalytic competence (Supplementary Fig. 11). In both the 70S IC-I and 70S IC* states, the catalytic His448 in switch II is oriented toward the nucleotide-binding pocket and positioned near the SRL nucleotide A2662, while both switch regions remain ordered (Fig. 3c, f; Supplementary Fig. 12a-c). This configuration is compatible with both pre-hydrolysis and immediate post-hydrolysis, pre–Pi-release states^32,33^, in which IF2α retains a GTP-like conformation. Because our cryo-EM analysis at this resolution cannot reliably distinguish between GTP and GDP–Pi, and because Pi retention can preserve a similar structural arrangement^32,33^ (Fig. 3f; Supplementary Fig. 11a), the 70S IC-I state likely represents a continuum of closely related nucleotide states. Within this continuum, some particles may retain GTP, while others have already undergone hydrolysis but still contain inorganic phosphate, resulting in nearly indistinguishable conformations.

Importantly, the structural conservation extends across the entire visible parts of IF2α. The N2-domain of IF2α remains anchored to helix h16 of the 16S rRNA, with density traceable from residue Ser294 onward (Fig. 3g). This interaction mirrors that observed in the 30S PICs (Figs. 2a, d) and in a homologous 70S IC structure from *P. aeruginosa*^26^ (Supplementary Fig. 10d), indicating that anchoring of the N2-domain to the 30S subunit is maintained after subunit joining. Despite this overall conservation in IF2α conformation, the most pronounced differences between 70S IC* and 70S IC-I emerge at the inter-subunit interface. Notably, H69 of the 23S rRNA in the 50S subunit of the 70S IC* state is curled back and does not engage with h44 of the 30S subunit as seen in 70S IC-I (Fig. 3h). Specifically, nucleotide A1913 in H69 is ∼13 Å apart from its position compared to 70S IC-I (Fig. 3h and Supplementary Fig. 8c). Thus, the presence of IF1 prevents the formation of inter-subunit bridge B2a. The neighbouring bridge, B2b, mediated through contacts between h24 in the 30S subunit with H68 in the 50S subunit, reaches only ∼14% of the buried surface area in 70S IC* compared to 70S IC-I (Supplementary Table 2 and Supplementary Fig. 8d). Furthermore, two inter-subunit bridges involving h23 and h24 remain incompletely established. Bridge B7a, which connects h23 of the 30S subunit to H68 of the 50S subunit, is only partially formed, reaching ∼40% of the buried surface area observed in 70S IC-I. In 70S IC*, the key h23 nucleotide A702 is displaced by ∼4 Å from its stacking interaction with A1848 of H68 (Fig. 3h, Supplementary Table 2 and Supplementary Fig. 8e), while bridge B7b, linking h23 and h24 to ribosomal protein uL2, is formed to only ∼32% of the buried surface area (Supplementary Table 2 and Supplementary Fig. 8f). These bridges, together with B2a, are among the key contacts required for stable 70S assembly^38^, with the establishment of B7a deemed to be a late event in 70S formation^39^. The partially formed inter-subunit interfaces in 70S IC* indicate that completion of the 70S IC requires IF1 dissociation. The markedly reduced contact area between the subunits in 70S IC*, by ∼1,603 Å^2^, is consistent with the proposed role of IF1 in promoting dissociation of vacant 70S and 70S–mRNA complexes^34^. In this state, IF1 interferes with the formation of bridges B2a, B2b, B7a and B7b, whereas IF2 compensates for this loss by contributing ∼2,000 Å^2^ of interaction surface with the 50S subunit, in agreement with its strong stimulation of subunit docking^5^. Together, these structures define a continuum of early 70S initiation states. The 70S IC* captures a pre-hydrolysis intermediate in which IF1 restricts subunit engagement, whereas 70S IC-I represents a more mature state following IF1 dissociation, in which inter-subunit bridges are fully established.

### Pi release and ribosome back-rotation drive IF2α compact conformation

The second initiation complex, 70S IC-II, provides further insights into the structural transitions that occur after GTP hydrolysis and prior to IF2α dissociation (Supplementary Table 1; Supplementary Figs. 4 and 9). This structure displays a less rotated state of the 30S subunit relative to the 50S subunit (3.4° compared to 4.1° in 70S IC-I) (Fig. 4a). Strikingly, IF2α adopts a compact conformation, reminiscent of the GDP-bound form previously observed only in *P. aeruginosa* 70S IC^26^ (Supplementary Fig. 10e-f). This configuration likely represents a late initiation intermediate, poised for IF2α release.

**Figure 4.**
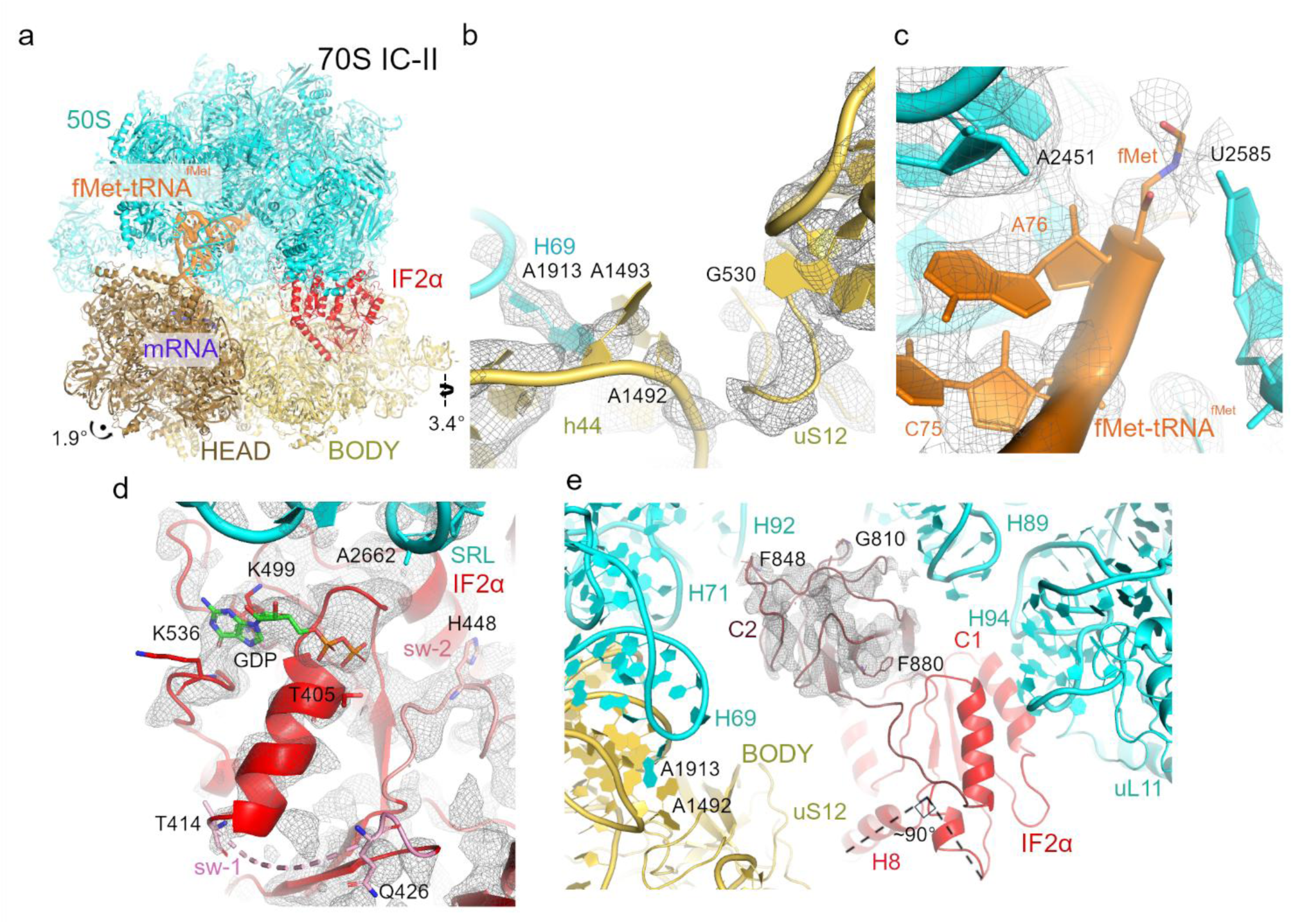
Cryo-EM structure of 70S IC-II complex. (**a**) Overall view of 70S IC-II, coloured as in Fig. 2. Arrows indicate the 30S head swivel and body rotation. (**b**) Close-up view of the DC, highlighting the positions of DC nucleotides A1492 and A1493. The original cryo-EM density map is shown as a grey mesh at 2.5 σ. (**c**) Close-up view into the PTC highlighting the 3′-CCA end of fMet-tRNAᶠᴹᵉᵗ. The fMet moiety and nearby 23S rRNA nucleotides are shown in sticks. The cryo-EM map for fMet-tRNAᶠᴹᵉᵗ and 23S rRNA was segmented (grey mesh) and contoured at 1.4 σ and 2.5 σ, respectively. (**d**) G-domain of IF2α in 70S IC-II. The purine base of GDP stacks against Lys499 and Lys536 while no density corresponding to the γ-phosphate is observed. Similarly, the switch I region (sw-1), spanning from Thr414 to Gln426, is not visible. Switch II (sw-2) is shown in salmon. The original cryo-EM map shown in grey mesh was segmented to IF2α G-domain region and SRL region and contoured at 2.0 σ and 3.5 σ, respectively. (**e**) Close-up view of the compact C2-domain of IF2α, highlighted in ruby. Selected residues are shown as sticks, while the rest of the IF2α structure, including the C1-domain and helix H8, are shown in red. The 90° kink of the C-terminal portion of helix H8 is displayed by dashed lines. The cryo-EM map was low-pass filtered (grey mesh; B-factor 50 Å^2^) and segmented corresponding to the C2-domain region and contoured at 1.5 σ.

Despite the dramatic rearrangement of IF2α, the DC remains flexible with reduced local resolution. The decoding nucleotides A1492 and A1493 are mirroring the configuration seen in 70S IC-I (Figs. 3e and 4b). In contrast to the preceding state, however, fMet-tRNA^fMet^ is no longer contacted by the IF2α C2-domain (Fig. 4a, c). Instead, it transitions to a P/P configuration (Fig. 5a). In this intermediate, the codon–anticodon base pairing is intact (Supplementary Fig. 5e), while the initiator tRNA elbow is shifted by ∼26 Å from its P/PI position in 70S IC-I (Fig. 5a). In addition, the fMet moiety is docked deeply within the peptidyl transferase centre (PTC) of 23S rRNA (Fig. 4c).

**Figure 5.**
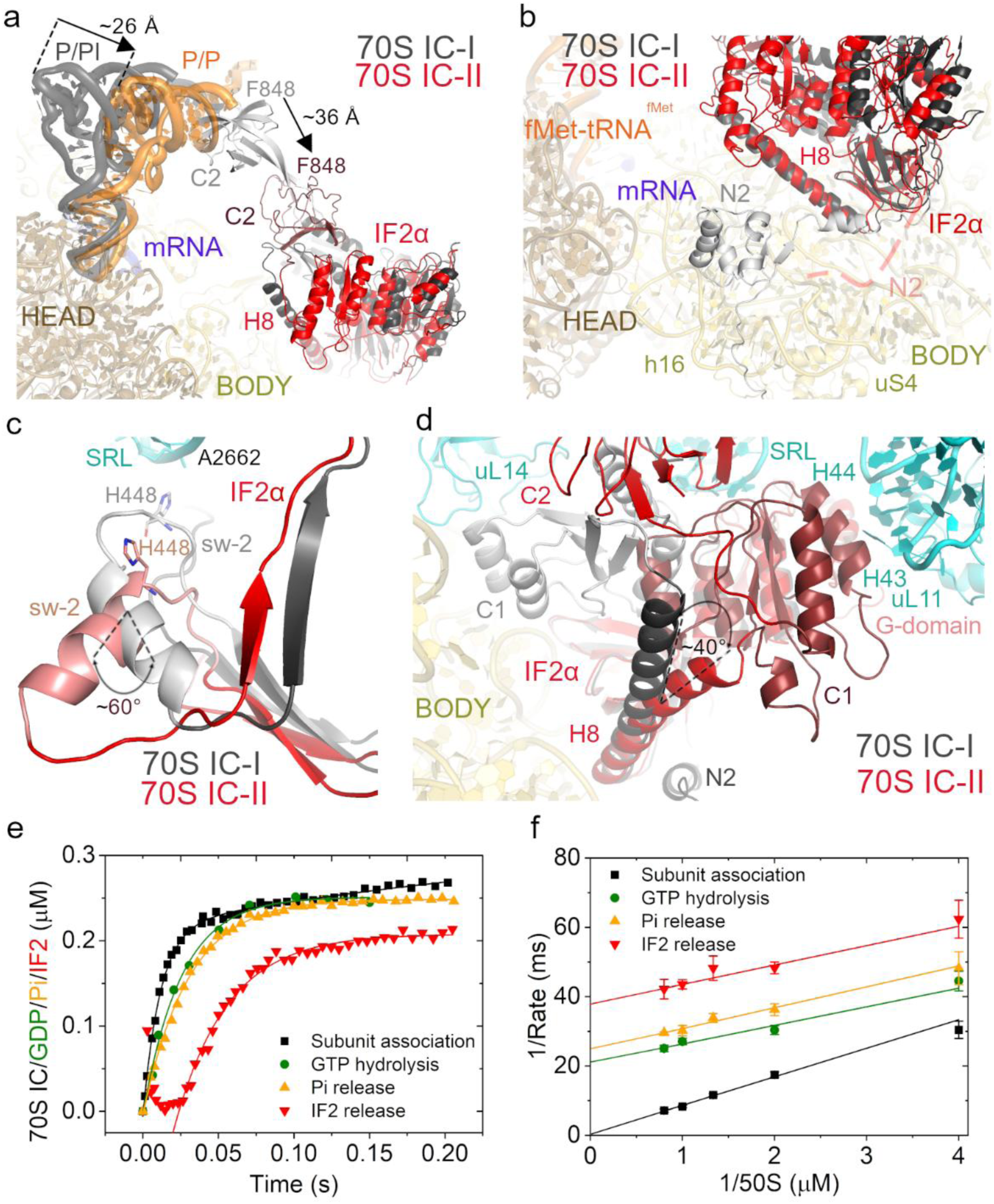
Transition between 70S IC-I and 70S IC-II and real-time monitoring of substeps of translation initiation. (**a**) Structural alignment of 70S IC-I and 70S IC-II depicting the transition of the fMet-tRNA^fMet^ from P/PI state (70S IC-I) to P/P state (70S IC-II). Distance between the elbow regions of fMet-tRNA^fMet^ is highlighted and depicted by an arrow. The retraction of the C2-domain is also highlighted (measured based on F848 position) and depicted by arrow. (**b**) Structural alignment depicting the absence of N2-domain of IF2α in 70S IC-II (dashed line) compared to 70S IC-I. (**c**) Structural alignment highlighting the repositioning of sw-2 region from 70S IC-I to 70S IC-II. The ∼60° rotation between sw-2 regions in both structures is depicted by dashed lines. (**d**) Structural alignment highlighting the displacement of C1-domain between 70S IC-I and 70S IC-II. The angle between H8 positions in both IF2α structures (∼40°) is depicted, along with selected helices of 23S rRNA. All shown structural alignments were performed by aligning the structures against 16S rRNA corresponding to the 30S body in 70S IC-II. The colour scheme for the 70S IC-II structure is as in Fig. 4, while the 70S IC-I structure is displayed using a domain-specific colour scheme in varying shades of grey. (**e**) Time courses of 70S IC formation defined by subunit association (black), GTP hydrolysis (green), Pi release (orange), and IF2α dissociation (red) as a function of time (representative plots). (**f**) Estimation of the time intervals between the substeps of translation initiation from the Lineweaver–Burk double reciprocal plots of the rates against 50S concentration (mean of three independent experiments with s.d.) at 37°C. At maximal 50S concentration, initial subunit association occurs very rapidly with mean time of ∼0.4 ms, followed by GTP hydrolysis at ∼21 ms, Pi release at ∼25 ms, and IF2α release at ∼38 ms. The experimental data are supplied in Supplementary Fig. 13.

Within the G-domain of IF2α, a clear GDP density is retained, indicating Pi release following GTP hydrolysis (Fig. 4d). Correspondingly, the sw-1 region (Thr414–Gln426), typically disordered after GTP hydrolysis^26,32,33^, is unresolved (Fig. 4d). In contrast, sw-2 remains well resolved but adopts a significantly different conformation (Figs. 4d, 5c), rotating by approximately 60°. This movement repositions His448 ∼14 Å away from the SRL compared to its location in 70S IC-I and results in tight stabilization of sw-2 through interactions with the C1-domain (Supplementary Fig. 12d, e). This reorganization marks the beginning of a structural rearrangement cascade triggered by GTP hydrolysis, with conformational changes propagating throughout IF2α. Such structural rearrangement is evident in the positioning of the C2-domain, which is now fully disengaged from the initiator tRNA. It retracts toward the ribosomal core, settling adjacent to 23S rRNA helices H71, H89, and H92 (Fig. 4e). This repositioning allows the C2-domain to form a compact, tightly associated interface with the C1 and G-domains (Supplementary Fig. 12d, e). Although the C2-domain exhibits considerable flexibility in this state, its position could still be determined, revealing the extent of its rearrangement.

Notably, the region defined by Phe848, previously involved in stabilizing the fMet moiety in 70S IC-I, is now displaced by ∼36 Å, highlighting the dramatic shift in domain reorganization (Fig. 5a). This prominent domain reorganization in IF2α continues through helix H8, with notable structural changes in its C-terminal segment. H8 is now redirected from its position in 70S IC-I by approximately 40° (Fig. 5d), with the C-terminal end of the segment forming a sharp 90° kink that allows it to engage directly with the C1-domain (Fig. 4e). The C1-domain is also displaced from its original position near the 30S subunit and relocates beneath the L11 stalk, now flanked by H95 of the 23S rRNA (Figs. 4e, 5d). In this new position, the remodeled sw-2 forms a contact with the C1-domain via helix H10 (Supplementary Fig. 12e), closely resembling the arrangement seen in the 70S initiation complex of *P. aeruginosa*^26^ (Fig. 5c and Supplementary Fig. 10e, f). Lastly, the propagation of conformational changes within IF2α together with continuous ribosomal back-rotation likely play a key role in promoting the detachment of the N2-domain. In 70S IC-II, the N2-domain is already released, contrasting its position in 70S IC-I (Fig. 5b). Here, only densities downstream of Ala397 are visible, indicating a loss of stable interaction. This progressive weakening of contacts after the Pi release suggests that IF2α is primed for dissociation.

Our fast-kinetics based biochemical data strongly support the structural observations of stepwise initiation complex remodelling. We have monitored kinetics of all substeps of translation initiation in the same experimental setup and estimated the mean catalytic times by titrating 50S to a fixed concentration of 30S PIC (Fig. 5e; Supplementary Fig. 13). Our results summarized in a Lineweaver–Burk double-reciprocal plot reveal a sequential timeline: initial subunit association occurs very rapidly with mean time of ∼0.4 ms, followed by GTP hydrolysis at ∼21 ms, Pi release at ∼25 ms, and IF2α release at ∼38 ms (Fig. 5f). The narrow 4 ms interval between the GTP hydrolysis and Pi release indicates that the GDP·Pi intermediate state is relatively short lived under our experimental conditions. Instead, a more pronounced time delay of ∼13 ms separates the Pi release event from the final dissociation of IF2α. This kinetic gap demonstrates that the primary rate limiting structural rearrangements required for IF2α dissociation occurs after GTP hydrolysis when Pi dissociates from the GTP pocket, which is in line with the prior report^29^.

Together, these biochemical and structural insights reinforce the idea that Pi release initiates a cascade of IF2α conformational changes. The resulting destabilization of IF2α contacts, particularly the detachment of its N2-domain and retraction of C2-domain, in combination with 30S back-rotation, likely drive IF2α release and facilitate the transition to a 70S EC ribosome.

Our final structure, termed as 70S EC, represents the endpoint of translation initiation (Supplementary Table 1; Supplementary Figs. 4, 9 and 14). Here, IF2α is no longer bound, and the ribosome adopts a non-rotated configuration^40,41^ (Supplementary Fig. 14a). The decoding centre is fully stabilized^11^ with clear density for decoding nucleotides A1492 and A1493 (Supplementary Fig. 14b). The fMet-tRNA^fMet^ is accommodated in the P/P position, exhibiting base pairing with the mRNA start codon (Supplementary Fig. 5f) and the fMet moiety is fully buried within the PTC (Supplementary Fig. 14c). This structure therefore reflects a fully matured 70S EC, marking the transition to an elongation-ready state.

## Discussion

Through a series of cryo-EM structures spanning 30S PICs to elongation-competent 70S ribosomes, combined with fast-kinetics measurements, we define key structural transitions during bacterial translation initiation in *E. coli* (Fig. 6). Rather than aiming to provide a comprehensive view of bacterial translation initiation – which would require the inclusion of all three initiation factors, including IF3 – our analysis focuses on the stages of initiation in the absence of IF3, revealing how IF1 and IF2 coordinate subunit joining and maturation of the 70S elongation complex. Our study highlights IF1’s role in interfering with the formation of inter-subunit bridges, consistent with previous biochemical studies^42,43^, while extending these observations by revealing its broader impact on multiple bridge interfaces. In doing so, IF1 shifts the burden of subunit docking onto IF2, thereby ensuring that only fully assembled 30S PICs join with 50S subunits^8,11^. This is consistent with earlier models proposing that the 30S-to-70S IC transition constitutes a late checkpoint for initiation fidelity^8,13^.

**Figure 6.**
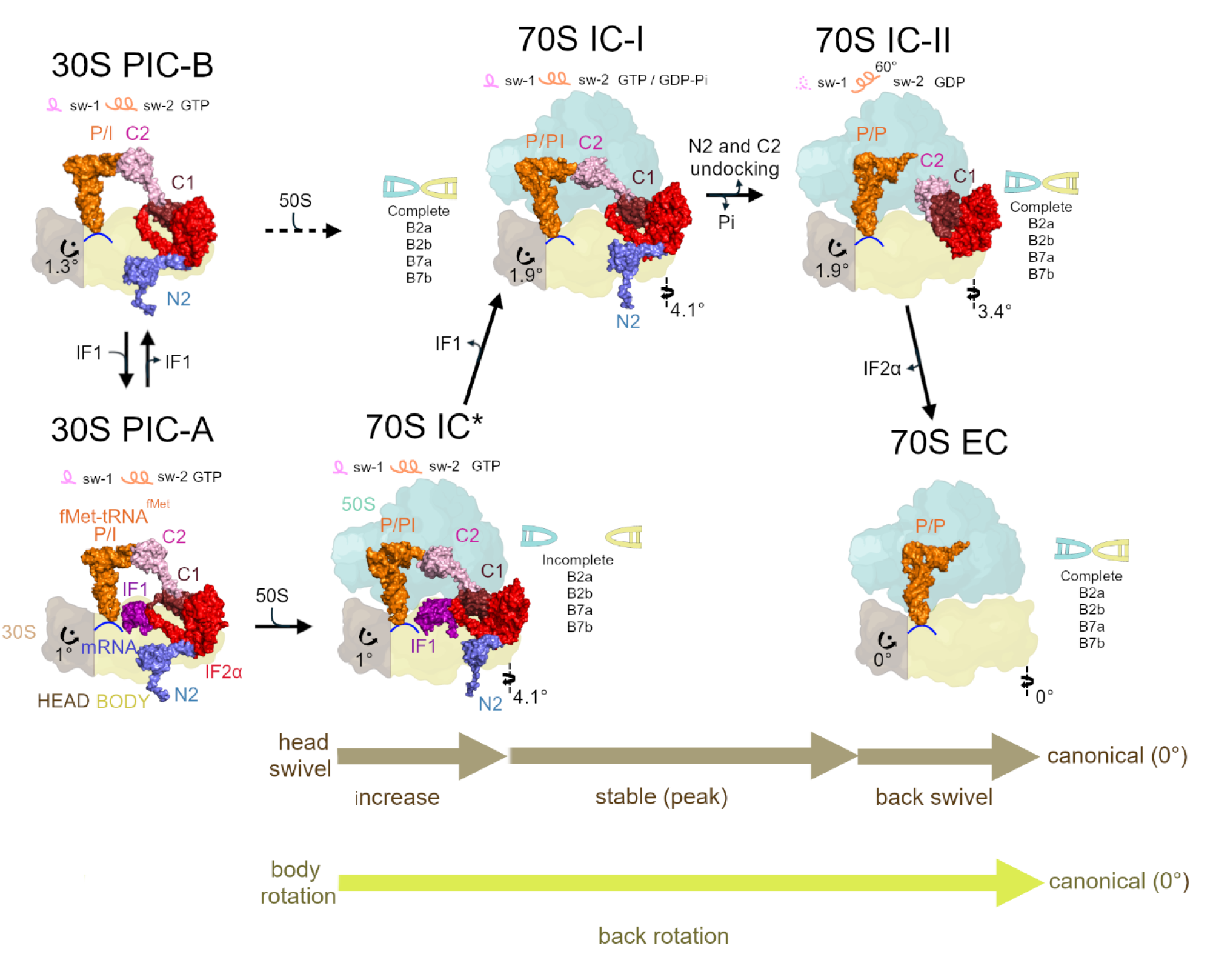
Model of translation initiation in bacteria. Illustration of the sequential conformational changes that occur during bacterial translation initiation, starting from the 30S PICs and progressing to 70S EC. This trajectory includes gradual head swiveling and body rotation, which reverse back to the canonical state in 70S EC. IF1 is retained in the pre-hydrolysis 70S IC*, where it impedes formation of inter-subunit bridges. Upon GTP hydrolysis, IF1 dissociates, allowing full 50S association. IF2α undergoes significant conformational rearrangements after Pi release, particularly in C2-, C1- and N2-domains, which remodel dynamically during the transition between 70S IC-I and II intermediates. The conformational states of switch 1 and switch 2 loops within the G-domain are also depicted. Ordered regions are shown as solid lines, while disordered loops appear as dashed lines. The presence of GDPCP mimicking the GTP-like state in 70S IC*, GTP/GDP-Pi and GDP are marked to reflect the stages of GTP hydrolysis and Pi release.

The coexistence of both IF1-bound and IF1-free 30S PICs (Fig. 2a, d) likely reflects a dynamic equilibrium in which IF1 enhances initiation fidelity without being strictly required for subunit joining and formation of a functional 70S IC^5,29^. More importantly, our data show that IF1 may persist in the 70S IC until GTP hydrolysis by IF2 (Fig. 3a), consistent with previous evidence for a late initiation intermediate containing both IF1 and IF2^8^. In the 70S IC* structure, IF1 perturbs subunit joining by inducing conformational changes in the 30S platform that disrupt the formation of inter-subunit bridges B2b, B7a and B7b (Supplementary Fig. 8d-f). Its position in the A site also sterically interferes with the central bridge B2a (Supplementary Fig. 8c). A previous 30S–IF1 crystal structure showed that IF1 alters the conformation of h44^44^, but this feature is absent in our IF1-bound 70S IC*, suggesting that the 30S subunit adopts a high-energy intermediate state with reduced IF1 affinity, as proposed earlier^34,42,43^. Rather than directly distorting h44, IF1 appears to stabilize its canonical conformation. The energetic consequences of IF1 binding may instead be accommodated through subtle rearrangements of the 30S platform domain, allowing conformational plasticity within the decoding centre without large-scale deformation of h44. These subtle rearrangements of the 30S platform domain can weaken inter-subunit bridges B2b, B7a and B7b. We propose that the reduced surface area at bridges B2a, B2b, B7a and B7b is offset by extensive IF2-50S interactions, which in turn strongly accelerate subunit docking^5^. Consistent with this interpretation, disruption of H69 in the 23S rRNA (in formation of B2a bridge) has been shown to impair late stages of initiation and formation of elongation-competent ribosomes^45,46^. This interplay between IF1-mediated destabilization of inter-subunit bridges and IF2-mediated stabilization of the subunit interface provides a structural basis for initiation quality control, consistent with previous models of IF2-dependent subunit docking^45,47^. It ensures that 30S PICs containing IF2–GTP in the extended conformation and fMet-tRNA^fMet^ in the P/PI state can form productive 70S ICs. Incorrectly assembled 70S ICs that lack initiator tRNA are rapidly destabilized by IF1 and IF3, consistent with a model in which IF1 promotes a transient, high-energy or partially open ribosome conformation that exposes the IF3 binding site and facilitates ribosome splitting^34^.

Moreover, our structures suggest that IF1 dissociation occurs after subunit joining but prior to IF2 release, consistent with previous studies^5,8,11,47^, although we cannot unambiguously distinguish whether this step is directly coupled to GTP hydrolysis or subsequent Pi release. In fact, dissociation of IF1 appears to coincide with the establishment of the inter-subunit bridge B2a, likely occurring during the transition from the 70S IC* to 70S IC-I (Fig. 3h). This model is supported by the position of IF1 in 30S PIC-A and 70S IC*, where its presence would sterically hinder B2a bridge formation (Fig. 3h). GTP hydrolysis by IF2α, together with the ribosomal back-rotation that accompanies the transition from the 70S IC* (∼4.1°) to the 70S EC (0°), drives the irreversible dissociation of IF1. This interplay underscores IF1 and IF2 as central regulators of the transition from the 30S IC to the 70S _IC_8,48.

IF2’s dual role as a structural scaffold and temporal regulator of translation initiation is further supported by visualization of the N-terminal N2-domain (Figs. 2h and 3g). This domain is positioned on the solvent-exposed side of the 30S subunit and remains associated across the 30S PICs, 70S IC* and 70S IC-I. These results, consistent with previous structural studies^26^, assign a major role for the N2-domain in stabilizing IF2 binding to the 30S subunit, essential for efficient subunit association during initiation. Notably, this domain is present in all IF2 isoforms and the tested IF2 variants (Fig. 1a). The similar rates of subunit association (Fig. 1b) with IF2α, D1 and D2 pre-bound to 30S PIC echo this notion, and moreover, suggest that the α-specific N-terminal extension (N1-domain) is not essential for rapid subunit association.

Our biochemical data in Fig. 1c, where IF2α, D1 and D2 variants show progressively slower rates of subunit association, at first appear inconsistent with the above conclusion. However, in this experiment, IF2 variants were supplied together with 50S subunits, and the overall reaction was rate-limited by IF2 binding to 30S PIC. Thus, the faster subunit association observed with IF2α compared with D1 and D2 (Fig. 1c) reflects more efficient binding of IF2α to the 30S PIC. Notably, the α-specific N-terminal extension remains unresolved in our structures, suggesting that its contribution is likely indirect and primarily important for the initial recruitment step rather than the later stages of initiation. Thus, our biochemical and structural data collectively confirm that the N1-domain of IF2α is important for efficient recruitment of IF2, while it plays no significant role in the subsequent steps of translation initiation. This interpretation is consistent with previous structural studies highlighting the role of the N2-domain in ribosome binding^20,35^.

The stepwise transition of IF2α from extended to compact form clarifies the role of GTP hydrolysis in coordinating translation initiation. In our 70S IC-I structure, IF2α remains in an extended conformation and tightly engaged with both the ribosome and the initiator tRNA. The active-site geometry of the G-domain is consistent with either a GTP-like or GDP–Pi-like configuration; however, these nucleotide states cannot be unambiguously distinguished at the current resolution. Similar active-site geometries have been observed in translational GTPases in GDP–Pi states^32,33^, supporting the interpretation that this state may represent a mixture of closely related nucleotide configurations. This structural ambiguity aligns with the biochemical data indicating that GTP hydrolysis alone does not trigger IF2 release but must be followed by Pi release^16,47^ (Fig. 5e, f). Notably, previous kinetic studies in different experimental setup have reported that Pi release can be slower than GTP hydrolysis^8,12,13^, suggesting that different physiological conditions may influence the observed lifetime of the GDP–Pi intermediate. Given the ∼13 ms delay between Pi release and IF2α dissociation from the ribosome (Fig. 5f), this interval is likely occupied by a relatively slow but substantial conformational rearrangement within IF2α. The structural reorganization of IF2 that follows Pi release is visualized in our 70S IC-II structure, where IF2α adopts a compact conformation (Fig. 4a, e). This conformation is characterized by the retraction of the C2-domain away from the initiator tRNA and PTC, displacement of the C1-domain toward the core of the 50S subunit and destabilization of the G-domain (Supplementary Fig. 12d). At the same time, the sw-2 region becomes reordered through interactions with the C1-domain, and the N2-domain loses contact with the 30S subunit (Fig. 5b, Supplementary Fig. 12e). Once Pi is released, IF2α must disengage from most of its extensive anchoring contacts with the ribosome before it can dissociate, a requirement that necessitates this substantial structural rearrangement. This behaviour mirrors the large conformational transitions undergone by other translational GTPases, such as EF-Tu and EF-G, during elongation^32,33^. The transitions between these IF2 intermediates also involve progressive 30S back-rotation, revealing coordinated factor remodelling and ribosomal resetting, indicating that IF2α compaction can occur before this rotation is complete.

Taken together, the structures presented in this study in combination with fast-kinetics converge into a model where initiation progresses through a series of tightly coupled rearrangements, each regulated by ribosomal dynamics, bridge remodelling, GTP-hydrolysis, and IF2α conformational changes (Fig. 6). Mapping this sequence of events refines current models of bacterial translation initiation and provides a structural framework for understanding late-stage subunit joining.

## Materials and Methods

### Cloning, expression and purification of the IFs

For cloning different variants of IF2 (IF2α, D1 -Δ1-159 and D2 -Δ1-289), corresponding segments of *E. coli infB* gene were PCR amplified and fused into the pET24a (Novagen) expression vector between NdeI and XhoI sites, which includes an N-terminal His6-tag separated by a TEV protease cleavage site. Successful clones were selected after confirmation by DNA sequencing. The *E. coli infA* gene encoding IF1 was also cloned in pET24a with uncleavable N-terminal His6-tag.

For over-expression of IFs, chemically competent *E. coli* BL21 (DE3) or *E. coli* BL21-CodonPlus (DE3)-RIPL cells (Agilent Technologies) cells were transformed with the individual IF clones. Expression and purification of the IFs were conducted using standard protocols^17,49,50^. Protein expression was induced by addition of IPTG (1 -1.5 mM) to the LB culture at 37°C at an OD600 of 0.4 -0.7. Cells were harvested after 3 -4 hrs by centrifugation at 4,500 rpm for 30 min, washed, and resuspended in buffer A (Supplementary Table 3). Cells were lysed by sonication in buffer A (kinetic studies) or by using a Microfluidics M-110P homogenizer (cryo-EM studies). The lysate was clarified by centrifugation at 15,000 -20,000 rpm for 1 h at 4 °C and the cleared lysate was loaded onto a 5 ml HisTrap Ni-NTA column (Cytiva) equilibrated with buffer A, using an ÄKTA Explorer / Pure FPLC system. After washing the desired IFs were eluted with linear gradient of buffer B, which is essentially buffer A supplemented with 200 – 500 mM imidazole (Supplementary Table 3).

For cryo-EM studies, affinity-purified IF1 and IF2α were dialyzed against buffer C (Supplementary Table 3) and further purified by ion-exchange chromatography using a HiPrep Q XL 16/10 column (Cytiva) equilibrated with the same buffer. IF2α was eluted with a linear gradient of buffer D (buffer C with 1 M NaCl), while IF1 was eluted in flow-through. Both proteins were dialyzed in respective buffer E (Supplementary Table 3) and then purified to homogeneity via size-exclusion chromatography using a HiLoad 16/600 Superdex 75 pg (for IF1) or 200 pg (for IF2α) column (Cytiva). Protein-containing fractions were pooled, buffer exchanged into respective storage buffer (Supplementary Table 3) and concentrated using a 3 kDa (IF1) or 30 kDa (IF2α) cut-off ultrafiltration unit (Millipore). The final purified proteins were flash-frozen in liquid nitrogen and stored at -80 °C.

For kinetic studies, the N-terminal His6-tag was removed from the IF2 variants by treatment with TEV protease in buffer A for 15 minutes at 30 °C. The TEV-treated proteins were reloaded onto the HisTrap Ni-NTA column, where the tag-free IF2 variants were eluted in flow-through in buffer A. The proteins were concentrated using Amicon® Ultra Centrifugal Filters and the concentrations were measured using Bradford assay. For Rhodamine labelling, IF2α was incubated in labelling buffer (20 mM Tris-HCl pH 7.2, 150 mM NaCl, 5 mM EDTA) with 10-fold excess of Tetramethylrhodamine-5-Maleimide (Thermo Fisher Scientific Inc.) for 16 hours at 4 °C. The reaction was stopped by adding βME. The residual free dye was removed by passing the labelling reaction through a HiTrap Q HP anion exchange column following previously published protocol^47,51^. IF2-Rho was eluted using high salt buffer (50 mM HEPES-KOH pH 7.5, 1 M NaCl, 10 mM MgCl2, 10% glycerol) and dialyzed in storage buffer (10 mM Tris-HCl pH 7.5, 50 mM NaCl, 5 mM MgCl2, 10% glycerol). The purified proteins were stored at -80 °C.

### Purification of 70S ribosomes and ribosomal subunits

Ribosomal subunits were purified from *E. coli* MRE600 cells as previously described^52^. Briefly, the cells were lysed and homogenized in a Microfluidics M-110P system in buffer A (20 mM Tris-HCl pH 7.0, 12.5 mM MgCl2, 100 mM NH4Cl, 0.5 mM EDTA, 6 mM βME) and the lysate was clarified by centrifugation at 15,000 rpm for 1 h at 4°C (JA-17 rotor). The supernatant was layered over a 37.7% sucrose cushion in buffer B (20 mM Tris-HCl pH 7.0, 15 mM MgCl2, 500 mM NH4Cl, 0.5 mM EDTA, 6 mM βME) and ultracentrifuged at 43,000 rpm for 16 h at 4°C (Ti45 rotor). The ribosomal pellet was resuspended in buffer A and centrifuged at 4,500 rpm for 5 min at 4°C. After adjusting the NH4Cl concentration to 500 mM in 40 ml buffer B, the sample was ultracentrifuged again at 55,000 rpm for 2 h at 4°C (Ti70 rotor). The resulting pellet was resuspended in 1 ml of buffer A and kept on ice. To isolate the 70S ribosomes, the sample was fractionated on a 10-35% sucrose gradient in buffer A (20,000 rpm, 13 h, 4°C; SW32 rotor). The 70S peak was collected and washed in buffer A (20 mM Tris-HCl pH 7.0,12.5 mM MgCl2, 100 mM NH4Cl, 0.5 mM EDTA, 6 mM βME). For the isolation of the 30S and 50S subunits, purified 70S ribosomes were incubated on ice for 30 minutes in buffer D (20 mM Tris-HCl pH 7.0,1.5 mM MgCl2, 500 mM NH4Cl, 0.5 mM EDTA, 6 mM βME). The sample was then loaded on 10%-35% sucrose gradient in buffer D and ultracentrifuged at 20,000 rpm for 13 h and 4°C (SW32 rotor). The 30S and 50S subunit fractions were collected, buffer exchanged and concentrated into buffer A. The ribosomal subunits in buffer A were flash-frozen in liquid nitrogen and stored at -80°C.

### fMet-tRNA^fMet^ and mRNA for cryo-EM

Initiator fMet-tRNA^fMet^ was over-expressed in a 1 L culture of exponentially growing *E. coli*. Several hours after reaching saturation, total tRNA (0.65 µmol) was isolated by differential centrifugation^53^ and deacylated for 3 h at 37 °C in 100 mM glycine buffer (pH 9.0). When required, initiator tRNA was affinity-purified by hybridization to a complementary biotinylated probe attached on streptavidin–sepharose^54^. After washing, bound tRNA was released at 65 °C for 5 min, ethanol-precipitated, and resuspended in water (yield ∼65%).

Charging and formylation proceeded in two steps. First, 40 nmol of affinity-purified tRNA was aminoacylated in a 100 µL reaction containing 10 µM MetRS, 1 mM methionine, 2 mM ATP-Mg, 0.2 mg/mL BSA, 50 mM Tris–HCl (pH 7.5), 20 mM KCl, 10 mM MgCl₂ and 4 mM DTT at 37 °C. After 10 min, charging efficiency was assessed by gel-shift analysis^55^. The reaction was then supplemented with 0.85 mM 10-formyltetrahydrofolate (generated from folinic acid at pH 8.0 for 15 min at RT) and 10 µM methionyl-tRNA formyltransferase. Stoichiometric formylation was achieved after 10 min at 37 °C. The reaction was quenched with 0.1 volume of 2.5 M sodium acetate (pH 5.0), extracted with phenol:chloroform:isoamyl alcohol (25:24:1), ethanol-precipitated, and resuspended in 25 mM sodium acetate (pH 5.0) for storage at −70 °C.

mRNAs containing the Shine-Dalgarno sequence and a linker to place the AUG codon in the P site were synthesized by IDT. The mRNA used for time-resolved cryo-EM data collection contains the sequence 5’-GCAA-<u>AGGAGG</u>-UAAAA-<u>AUG</u>-GUU-ACA-CUC-AAG-AUU-GCC-AUC-AUU-GAC-UUA-GCC-GGA-CA-3’. The mRNA used for trapped 70S IC* complex with GDPCP contains the sequence 5’-GGCA-<u>AGGAGG</u>-UAAAA-<u>AUG</u>-UUC-UAA-3’. The lyophilized mRNAs were resuspended in Milli-Q water, adjusted to a final concentration of 100 µM and stored at −20 °C until use.

### Translation initiation kinetics and analysis

All kinetics experiments were carried out in HEPES-polymix buffer^49^ (5 mM HEPES (pH 7.5), 5 mM NH4Cl, 5 mM Mg(OAc)2, 100 mM KCl, 0.5 mM CaCl2, 8 mM putrescine, 1 mM spermidine and 1 mM dithioerythritol (DTE)) supplemented with 1 mM ATP and 1 mM GTP. The 70S ribosome and the 50S and 30S subunits, XR7 Met-Leu-Leu mRNA, BOP-Met-tRNA^fMet^ and [^3^H]fMet-tRNA^fMet^ were purified as described^49,56^. All translation components (ribosome, subunits, initiation factors and tRNAs) were either purified from or recombinantly expressed in *E. coli*.

To test the IF2 variants in subunit association assay, a 30S PIC containing 30S (0.5 μM), XR7 Met-Leu-Leu mRNA (2 μM), IF1 (2 μM) and fMet-tRNA^fMet^ (2 μM) was pre-formed by incubation at 37 °C for 10 min. The 50S (0.5 μM) was also incubated separately. The IF2 variants (IF2α, D1 and D2) (2 μM) were added either to the 30S PIC or to the 50S mix. Subunit association was initiated by rapid mixing of the 30S PIC and 50S in stopped-flow (Biologic) and the time course of association was followed by increase in Rayleigh light scattering at 365 nm. The experiments were repeated also in the presence of IF3 (2 μM).

For complete kinetic analysis of all substeps of translation initiation, subunit association, corresponding GTP hydrolysis, Pi release and IF2 release experiments were conducted following previously published protocols^47^ with minor modifications. In all experiments, a preformed 30S PIC mix containing 30S (0.5 μM), XR7 Met-Leu-Leu mRNA (2 μM), IF1 (2 μM), IF2 variant (2 μM), and fMet-tRNA^fMet^ (2 μM), was rapidly mixed in stopped-flow or quench-flow with 50S (0.5 μM). Briefly, subunit association was followed by increase in Rayleigh light scattering (365 nm) in stopped-flow, GTP hydrolysis was measured by using ^3^H-GTP in quench-flow followed by HPLC analysis of ^3^H-GTP/GDP, Pi release was monitored in stopped-flow by fluorescence increase at 425 nm due to binding of Pi to MDCC-labeled phosphate binding protein (PBP), and IF2-Rhodamine release was followed by the decrease in rhodamine fluorescence at 590 nm (λEx = 560 nm) in stopped-flow. Pi release and IF2-Rho release were followed simultaneously with subunit association, using dual detectors in an Applied Photophysics stopped-flow. Binding of fMet-tRNA^fMet^ to the 70S IC was also followed using BOP-Met-tRNA^fMet^ in stopped-flow, by monitoring BOP fluorescence at 590 nm (λEx = 560 nm) as published earlier^40^. All kinetic data are fitted with OriginPro 2016 (Origin LabCorp) and rates ± s.d. are estimated from three independent biological repeats.

### Trapped 70 IC* cryo-EM grid preparation, data collection and image processing

The 30S pre-initiation complex (30S PIC) was reconstituted by mixing *E. coli* MRE600 30S ribosomal subunits, *E. coli* IF1, IF2α, and initiator fMet-tRNA^fMet^. Briefly, the complex was assembled by mixing components to a final concentration of 8 µM 30S, 8 μM mRNA and 10 μM fMet-tRNA^fMet^ in the association buffer containing 10 mM HEPES-KOH pH 7.5, 60 mM KCl, 15 mM NH4Cl, 10 mM MgCl2 and 6 mM βME, incubated at 37 °C for 10 minutes and brought to room temperature. Initiation factors and nucleotide corresponding to a final concentration of 50 μM IF1, 30 μM IF2α, and 1 mM GDPCP (Sigma) were added to the 30S, tRNA and mRNA mixture and incubated at room temperature for 10 minutes. The 70S initiation complex was assembled with the addition of 1.9 µM *E. coli* 50S ribosomes (final concentration) in the association buffer and incubated at 37°C for 5 minutes and transferred to ice. Quantifoil R2/1 Au 200 mesh grids (Electron Microscopy Sciences, Hatfield, PA) were pre-cleaned for 50 seconds in a Solarus 950 plasma cleaner (Gatan) and immediately used for freezing in the Leica EM GP2 cryo-plunger. 4 µL of the 70S initiation complex sample was applied to the grids and blotted for 3 seconds in 22°C, 85% humidity before plunging into liquid ethane. Grids were then transferred into a Titan Krios G3i electron microscope (Thermo Fisher Scientific) operating at 300 keV and data acquired with a K3 direct electron detector (Gatan) in counting mode, coupled with a BioQuantum electron energy filter (Gatan) using a slit width of 20 eV. The image stacks (movies) were acquired with pixel size of 0.8623 Å/pixel using the SerialEM software to record movies with 40 fractions and total accumulated dose of 40.299 e^-^/Å^2^/movie. A total of 12,290 micrographs were collected with defocus values ranging from –0.7 to –2.0 µm.

Data processing was done in CryoSPARC 4.7.1^57^. The image stacks were motion-corrected with the patch motion correction job, followed by patch contrast transfer function (CTF) estimation. Based on relative ice thickness, CTF fit, and total frame motion, 11,759 micrographs were selected for further processing. 2,996,938 particles were filtered based on defocus adjusted power and pick scores, extracted with a box size of 512 x 512 pixels, and subjected to three rounds of reference-free two-dimensional (2D) classification. Selected 2D classes containing 757,723 particles were used in an ‘ab-initio reconstruction’ job to generate three 3D volumes that were further used for ‘heterogeneous refinement’. One main class average of 70S ribosomes containing 530,976 particles was obtained, in addition to one 30S class containing 115,434 particles and one junk class that was discarded (Supplementary Fig. 7). The 30S ribosome class was classified using a 3D classification job resulting in 29,113 particles of 30S ribosomes with strong density for IF1, P-site fMet-tRNA^fMet^ and weak density for IF2α and a 70S IC class of 80,840 particles. These lowly populated classes were not processed further. The 70S ribosome class was bound to IF2α in the extended conformation and P-site fMet-tRNA^fMet^, which was classified using a soft mask on IF2α and the tRNA using a focused 3D classification job resulting in 447,736 particles of 70S ribosomes bound to IF1 and IF2α corresponding to 84.3% of the total number of 70S IC particles, a class of 40,523 particles of 70S ribosomes without IF1 and bound to compact IF2α, and a class of 42,716 particles of 70S ribosomes with a swivelled head, bound to IF2α and P/E-tRNA. The main 70S ribosome class bound to IF1 and IF2α was further classified using a focused 3D classification job followed by a focused 3D variability job with a mask on IF1, resulting in a class of 64,469 particles of 70S ribosomes bound to IF1 and IF2α (70S IC* state) and a class of 382,672 particles bound to IF2α only, corresponding to 12.3% and 72% of the total 70S IC particles, respectively. Non-uniform refinement with CTF refinement, per-particle defocus optimization performed on the class of 70S ribosomes bound to IF1 and IF2α (70S IC*) resulted in a final reconstruction with a resolution of 3.0 Å according to the 0.143 FSC criterion and sharpened with a B-factor of –52.6 Å^2^ from a final set of 64,469 particles, as detailed in Supplementary Table 1 and Supplementary Fig. 7. Local-resolution filtering was applied using blocres and blocfilt from the Bsoft package (v1.9.1)^58^.

### Ensemble cryo-EM grid preparation, data collection and image processing

Reactions were carried out in reassociation (RA) buffer (20 mM Tris-HCl pH 7.5, 50 mM KCl, 10 mM Mg(OAc)2, 4 mM βME). The 30S subunit mixture was assembled in a total volume of 20 µL. First, 2 µM 30S subunits were pre-activated by incubation for 5 min at 42°C in RA buffer. The mixture was then supplemented with 3 µM mRNA and 6 µM IF1 (final concentrations), incubated for 10 min at 37°C and subsequently placed on ice. In parallel, the 50S subunit mixture was prepared in a separate 20 µL volume containing 2 µM 50S subunits, 6 µM IF2α, 4 µM fMet-tRNA^fMet^, and 1 mM GTP (final concentrations). This mixture was incubated for 5 minutes at room temperature and then placed on ice. In both reactions, the molar ratio of ribosomal subunits to initiation factors was maintained at 1:3. To assemble the final initiation complexes, 8 µL from each of the 30S and 50S mixtures were rapidly combined on ice, yielding a final concentration of 1 µM ribosomal complexes. Immediately, 3 µL of the final mixture was applied to glow-discharged Quantifoil R2/1 300 mesh grids (glow-discharged for 30-40 s at 40 W ± 5 W using a Gatan Solarus II). The grids were blotted at ∼5 °C and ∼95% humidity and plunge-frozen into liquid ethane using an FEI Vitrobot Mark IV (Thermo Fisher Scientific). The time between mixing the two reactions and plunge-freezing was ∼20 seconds.

Two datasets were collected from the same cryo-EM grid containing the initiation reaction. In total, 17,477 micrographs were recorded using a Titan Krios transmission electron microscope operating at 300 kV (Thermo Fisher Scientific). Data were acquired with a K3 direct electron detector in counting mode, coupled with a BioQuantum energy filter (Gatan Inc.) using a slit width of 10 eV. The data acquisition was carried out using SerialEM^59^, with a defocus range of -0.5 to -2.1 µm. Each exposure was recorded as a movie consisting of 40 frames over a 2 s duration, with a total electron dose of 40 e⁻/Å^2^ per movie. The nominal magnification was 105,000×, yielding a pixel size of 0.834 Å. The movies were motion-corrected using MotionCorr2^60^. Defocus values for each micrograph were determined using cisTEM (v1.0-beta)^61^ with CTFFIND4^62^ integrated within the cisTEM interface. After inspecting the power spectra, 1,028 poor-quality movies were excluded from the original dataset. Particle picking in cisTEM yielded a total of 1,889,196 particles, which were subjected to 2D classification into 50 classes. From these, classes corresponding to 30S and 70S ribosomal particles, containing 351,980 and 825,112 particles, respectively, were selected for further processing. Unbinned particle stacks for both 30S and 70S datasets (box size of 480 pixels) were exported from cisTEM. Additionally, 8×, 4×, and 2× binned versions of each particle stack were generated using the resample.exe utility from the Frealign package^63^.

Particle alignment, 3D classification, and final refinements were performed using Frealign^63^. The 30S and 70S particle stacks were processed following the same alignment strategy. As initial, non-biased references, 30S and 70S ribosome structures without any factors from PDB 6WDD^32^ were converted to a map via EMAN2.1 package^64^ and resampled to a pixel size of 6.672 Å (matching the 8× binned particle stacks) and low-pass filtered to 30 Å resolution. These filtered volumes were used as starting references for global alignment. The alignment was initially performed on the 8× binned particle stacks, using C1 symmetry and Frealign mode 3 (global search) over 5 cycles in the resolution range of 300–30 Å. This was followed by a local refinement in mode 1, also for 5 cycles over the same resolution range, using the references obtained in the previous step. All subsequent alignment steps were carried out in mode 1. Next, the 4× binned stacks were aligned to the common reference using a stepwise increase in resolution limit (30 Å → 18 Å → 12 Å), with 5 cycles per resolution step. Finally, the 2× binned stacks were refined in the same way, with resolution steps of 12 Å → 10 Å → 8 Å. For each particle stack, 3D map reconstructions were generated using the top 60% of particles based on alignment scores. These reconstructions were visually inspected using Chimera^65^. The final alignment parameters from the 2× binned stacks were then used for 3D classification of both the 30S and 70S particle sets.

The aligned 2× binned 30S particle stack (351,980 particles) was first subjected to 3D classification into 10 classes at 8 Å resolution over 50 cycles. This initial classification revealed one high-resolution class representing 30S subunit bound to IF2α, alongside several low-resolution or empty/unbound 30S classes and one mixed class containing both 30S and 70S particles. From the high-resolution IF2α-containing 30S class, 51,248 particles were extracted (with minimal 50% occupancy) using the merge_classes.exe utility from the Frealign package^63^. This particle subset was then further 3D classified into two classes (8 Å resolution over 50 cycles) using a spherical focused mask with a 30 Å radius positioned to cover both the A-site, which corresponds to the IF1 binding region, and helix 8 of IF2α. This focused 3D sub-classification resulted in two distinct populations: one class of 22,197 particles that retained IF1, and another of 28,862 particles in which IF1 was absent. Both particle sets were extracted as previously described and subjected to an additional round of focused 3D classification into two classes, using a 40 Å spherical mask centred on the IF2α C2-domain and the fMet-tRNA^fMet^, to investigate potential structural heterogeneity in this region. While this final classification highlighted conformational flexibility in the IF2α C2-domain and the fMet-tRNA^fMet^ elbow, as illustrated in Supplementary Fig. 6, only minimal differences were observed in the remainder of the 30S complexes. Consequently, both main particle classes, one representing the 30S initiation complex containing both IF1 and IF2α together with fMet-tRNA^fMet^ positioned in PI state (designated 30S PIC-A), and the other representing the complex with IF2α alone with fMet-tRNA^fMet^ positioned in PI state (30S PIC-B), were used directly for final refinement. The final refinement of the corresponding unbinned particle stacks was performed in mode 1 over 5 cycles, and the final 3D reconstructions were generated using 95% of particles with the highest scores. This procedure yielded final high-resolution maps for both 30S PIC-A (22,197 particles) and 30S PIC-B (28,862 particles) at ∼3.0 Å resolution, based on the 0.143 FSC criterion, as reported in Supplementary Table 1 and Supplementary Fig. 3.

The aligned 2× binned 70S particle stack, consisting of 825,112 particles, was first subjected to 3D classification into 20 classes at 8 Å resolution over 50 cycles. This initial classification yielded 13 high-resolution 70S classes, 4 low-resolution 70S classes, and 3 classes corresponding to isolated 50S subunits. The 13 high-resolution 70S classes, comprising 614,621 particles, were (with minimal 50% occupancy) extracted using the merge_classes.exe utility from the Frealign package^63^. This sub-stack was then further classified into 14 classes (8 Å resolution over 50 cycles) using a spherical focused mask with a 40 Å radius centred on the IF2α binding site of the 70S ribosome. This focused classification revealed three major populations. One class, containing 26,032 particles, showed clear density for fMet-tRNA^fMet^ in the P/PI state and IF2α in its classical extended conformation. A second group, comprising 6 classes and totalling 214,068 particles, displayed fMet-tRNA^fMet^ in a P/E hybrid state along with density for the core region of IF2α in a more compact conformation. The remaining 7 classes, encompassing 324,406 particles, showed no detectable density for IF2α and a mixture of fMet-tRNA^fMet^ in P/E hybrid and P/P states, and were interpreted as post-initiation 70S complexes.

The extracted sub-stack corresponding to 70S complex with IF2α in its extended conformation was further classified into 3 classes (8 Å resolution over 50 cycles) using a spherical focused mask with a 35 Å radius positioned to encompass the IF2α C2-domain and the 3′-CCA end of the fMet-tRNA^fMet^. This classification aimed to better resolve the 70S state in which IF2α adopts the classical extended conformation. As a result, one high-resolution class containing 12,354 particles was obtained, representing the 70S initiation complex with fMet-tRNA^fMet^ in the P/PI state and IF2α in its fully extended form, with GTP bound in the G-domain. This complex was designated as 70S IC-I.

The sub-stack representing IF2α in a more compact conformation was next classified into 10 classes (8 Å resolution over 50 cycles), using a 35 Å spherical mask positioned on the ribosomal P-site. The goal was to distinguish non-enzymatic 70S complexes containing fMet-tRNA^fMet^ in the hybrid P/E state from true enzymatic complexes with the tRNA in the P/P state. Among these, only one class, consisting of 25,393 particles, corresponded to a 70S initiation complex with fMet–tRNA^fMet^ in the P/P state and partial occupancy of IF2α in a compact conformation. This particle stack was re-extracted and subjected to a final round of 3D classification into 3 classes (8 Å resolution over 50 cycles), using a 35 Å focused mask centered on the IF2α binding region. This classification identified one high-resolution class of 8,740 particles in which the C2-domain of IF2α was fully retracted, confirming the compact conformation. In this complex, termed as 70S IC-II, IF2α contains GDP in its G-domain and the fMet–tRNA^fMet^ was found in the P/P state.

The particle sub-stack representing post-initiation 70S complexes lacking IF2α was also subjected to further classification into 10 classes (8 Å resolution over 50 cycles), again using a 35 Å focused mask on the ribosomal P-site to identify elongation-competent 70S states. One high-resolution class of 31,752 particles showed fMet-tRNA^fMet^ stably bound in the canonical P/P state, representing the elongation-competent 70S complex, designated as 70S EC.

Final refinements of the corresponding unbinned particle stacks for the 70S IC-I, 70S IC-II, and 70S EC states were carried out using mode 1 over 5 cycles. For each, 95% of particles with the highest scores were used to generate the final 3D reconstructions. This process yielded high-resolution cryo-EM maps at ∼3.1 Å for 70S IC-I (12,354 particles), ∼3.1 Å for 70S IC-II (8,740 particles), and ∼2.9 Å for 70S EC (31,752 particles), according to the 0.143 FSC criterion, as detailed in Supplementary Table 1 and Supplementary Fig. 9.

Together with the previously refined 30S PIC-A and 30S PIC-B structures, all five final cryo-EM maps were used for atomic model building and structural refinement. Local-resolution filtering was applied using blocres and blocfilt from the Bsoft package (v1.9.1)^58^, and sharpening was performed with bfactor.exe from the Frealign package^63^ using B-factors in the range of −20 to −40 Å^2^ to enhance high-resolution map features. The final Fourier Shell Correlation (FSC) curves were computed between half-maps derived from even and odd particle sets using Frealign.

### Model building and refinement

Cryo-EM structures PDB 6WDD^32^, 8G7P^66^ and 7UNV^26^ were used as a starting model for structure refinements. The models of IF1 and IF2α in its extended conformation were generated using AlphaFold^67^ based on *E. coli* sequences using template structures from PDB 5LMV^10^ and PDB 7UNV^26^, respectively. The compact conformation was obtained by extracting IF2α from PDB entry 7UNQ^26^ and submitting it to D-I-TASSER (https://zhanggroup.org//D-I-TASSER/), using the *E. coli* IF2α sequence to guide the modelling. The structural models were initially rigid-body fitted into the corresponding cryo-EM maps using Chimera^65^. The manual modelling was performed using Coot^68^. Poorly defined regions within the cryo-EM maps such as IF2α loops or switch regions were omitted from the models when necessary. All structures were refined using phenix.real_space_refine in Phenix^69^. Secondary-structure and base-pairing restraints were applied to rRNA and tRNA, along with restraints for the ester bond between the 3′-CCA end of tRNA^fMet^ and the methionine residue, as well as for GTP or GDP, when necessary. Correlation coefficients reflecting the model-to-map fit in Phenix were carefully monitored throughout to prevent overfitting. The refined structural models showed strong agreement with the corresponding cryo-EM maps, as indicated by high correlation values. FSC values between the final models and maps were computed in Phenix using phenix.mtriage showing strong agreement between the structural models and cryo-EM maps. All resulting models have favourable stereochemical parameters, including minimal deviation from ideal bond lengths and angles, and a low number of macromolecular backbone outliers, as detailed in Supplementary Table 1. Structure quality was assessed using MolProbity^70^ and comprehensive cryo-EM validation tool in Phenix 1.19.2^71^. All structure superpositions, distance and rotation calculations and figure generation were performed using Chimera^65^ and PyMOL (The PyMOL Molecular Graphics System, Version 2.3.1, Schrödinger, LLC).

### Inter-subunit bridge buried surface area (BSA) calculations

Buried surface area (BSA) measurements were performed using PISA (https://www.ebi.ac.uk/pdbe/pisa/). BSA for 70S IC* (bound to IF1) and 70S IC-I (without IF1) were calculated and pairwise comparisons between the structures are tabulated as BSA loss, % loss, and % retained (Supplementary Table 2).

## Acknowledgements

This study was supported and funded by Czech Science Foundation, project no. 26-20643S (to G.D.), by the Swedish Research Council [2018-05946, 2018-05498 and 2023-05237]; and Wenner-Gren Foundation [UPD2023-0185, UPD2017-0238 and UPD2018-0306] (to S.S.); by the National Institute of General Medical Sciences of the National Institutes of Health grants R01-GM136936 and R35-GM158272 (to M.G.G.), the Welch Foundation grant H-2032-20230405 (to M.G.G.), by the NIH grant R35-GM134931 (to Y.M.H.), and by a training fellowship from the Gulf Coast Consortia, on the Houston Area Molecular Biophysics Program (NIH grant T32-GM150582 to N.A.V.). We gratefully acknowledge the Cryo-electron microscopy and tomography core facility CEITEC MU of CIISB, Instruct-CZ Centre, supported by MEYS CR (LM2023042) and the European Regional Development Fund-Project Innovation of Czech Infrastructure for Integrative Structural Biology (No. CZ.02.01.01/00/23_015/0008175). Computational resources were provided by the e-INFRA CZ project (ID:90254), supported by MEYS CR. We thank Dr. Michael B. Sherman for advice and support with cryo-EM data collection, Drs. Ka-Yiu (Clem) Wong and John Perkyns for computational support, and the Sealy and Smith Foundation for supporting the Sealy Center for Structural Biology and Molecular Biophysics at the University of Texas Medical Branch. Open access charge is covered by Uppsala University library. The funders had no role in study design, data collection and analysis, decision to publish, or manuscript preparation.

## Author Contributions

Conceptualization: M.G.G., S. S., G.D. Resources: G.S.G, H.Z., X.G., R.S.B., C.H., A.H., N.A.V., S.B., L.S., C.S.M., H.G., Y.M.H., M.G.G., S.S., G.D. Biochemistry experiments and analysis: C.H., C.S.M., X.G. and S.S.; cryo-EM experiments and structure determination: G.S.G., H.Z., R.S.B., A.H., N.A.V., M.G.G., G.D.; Writing-Original Draft G.S.G, M.G.G., S.S. and G.D.; Writing-Review and Editing: G.S.G, H.Z., X.G., R.S.B., C.H., A.H., N.A.V., S.B., H.G., Y.M.H., M.G.G., S.S., G.D.; Supervision: M.G.G., S. S., G.D. Funding acquisition: Y.M.H., M.G.G., S.S. and G.D.

## References

1. Milon, P. & Rodnina, M.V. Kinetic control of translation initiation in bacteria. Crit Rev Biochem Mol Biol 47, 334–48 (2012).

2. Gualerzi, C.O. & Pon, C.L. Initiation of mRNA translation in bacteria: structural and dynamic aspects. Cell Mol Life Sci 72, 4341–67 (2015).

3. Samatova, E., Daberger, J., Liutkute, M. & Rodnina, M.V. Translational Control by Ribosome Pausing in Bacteria: How a Non-uniform Pace of Translation Affects Protein Production and Folding. Front Microbiol 11, 619430 (2020).

4. Rodnina, M.V. Translation in Prokaryotes. Cold Spring Harb Perspect Biol 10(2018).

5. Antoun, A., Pavlov, M.Y., Lovmar, M. & Ehrenberg, M. How initiation factors tune the rate of initiation of protein synthesis in bacteria. EMBO J 25, 2539–50 (2006).

6. Dahlquist, K.D. & Puglisi, J.D. Interaction of translation initiation factor IF1 with the E. coli ribosomal A site. J Mol Biol 299, 1–15 (2000).

7. Fabbretti, A. et al. The real-time path of translation factor IF3 onto and off the ribosome. Mol Cell 25, 285–96 (2007).

8. Goyal, A., Belardinelli, R., Maracci, C., Milon, P. & Rodnina, M.V. Directional transition from initiation to elongation in bacterial translation. Nucleic Acids Res 43, 10700–12 (2015).

9. Julian, P. et al. The Cryo-EM structure of a complete 30S translation initiation complex from Escherichia coli. PLoS Biol 9, e1001095 (2011).

10. Hussain, T., Llacer, J.L., Wimberly, B.T., Kieft, J.S. & Ramakrishnan, V. Large-Scale Movements of IF3 and tRNA during Bacterial Translation Initiation. Cell 167, 133–144 e13 (2016).

11. Kaledhonkar, S. et al. Late steps in bacterial translation initiation visualized using time-resolved cryo-EM. Nature 570, 400–404 (2019).

12. Milon, P., et al. Transient kinetics, fluorescence, and FRET in studies of initiation of translation in bacteria. Methods Enzymol 430, 1–30 (2007).

13. Milon, P., Konevega, A.L., Gualerzi, C.O. & Rodnina, M.V. Kinetic checkpoint at a late step in translation initiation. Mol Cell 30, 712–20 (2008).

14. Tsai, A. et al. Heterogeneous pathways and timing of factor departure during translation initiation. Nature 487, 390–3 (2012).

15. Milon, P. et al. The ribosome-bound initiation factor 2 recruits initiator tRNA to the 30S initiation complex. EMBO Rep 11, 312–6 (2010).

16. Tomsic, J. et al. Late events of translation initiation in bacteria: a kinetic analysis. EMBO J 19, 2127–36 (2000).

17. Antoun, A., Pavlov, M.Y., Tenson, T. & Ehrenberg, M.M. Ribosome formation from subunits studied by stopped-flow and Rayleigh light scattering. Biol Proced Online 6, 35–54 (2004).

18. Plumbridge, J.A. et al. Cloning and mapping of a gene for translational initiation factor IF2 in Escherichia coli. Proc Natl Acad Sci U S A 79, 5033–7 (1982).

19. Cenatiempo, Y. et al. The protein synthesis initiation factor 2 G-domain. Study of a functionally active C-terminal 65-kilodalton fragment of IF2 from Escherichia coli. Biochemistry 26, 5070–6 (1987).

20. Simonetti, A. et al. Involvement of protein IF2 N domain in ribosomal subunit joining revealed from architecture and function of the full-length initiation factor. Proc Natl Acad Sci U S A 110, 15656–61 (2013).

21. Brandi, A. et al. Transcriptional and post-transcriptional events trigger de novo infB expression in cold stressed Escherichia coli. Nucleic Acids Res 47, 4638–4651 (2019).

22. Brandi, A. et al. Translation initiation factor IF2 contributes to ribosome assembly and maturation during cold adaptation. Nucleic Acids Res 47, 4652–4662 (2019).

23. Laursen, B.S., Mortensen, K.K., Sperling-Petersen, H.U. & Hoffman, D.W. A conserved structural motif at the N terminus of bacterial translation initiation factor IF2. J Biol Chem 278, 16320–8 (2003).

24. Laursen, B.S., Kjaergaard, A.C., Mortensen, K.K., Hoffman, D.W. & Sperling-Petersen, H.U. The N-terminal domain (IF2N) of bacterial translation initiation factor IF2 is connected to the conserved C-terminal domains by a flexible linker. Protein Sci 13, 230–9 (2004).

25. Caserta, E. et al. Translation initiation factor IF2 interacts with the 30 S ribosomal subunit via two separate binding sites. J Mol Biol 362, 787–99 (2006).

26. Basu, R.S., Sherman, M.B. & Gagnon, M.G. Compact IF2 allows initiator tRNA accommodation into the P site and gates the ribosome to elongation. Nat Commun 13, 3388 (2022).

27. Caserta, E. et al. Ribosomal interaction of Bacillus stearothermophilus translation initiation factor IF2: characterization of the active sites. J Mol Biol 396, 118–29 (2010).

28. Metelev, M. & Johansson, M. A complex between IF2 and NusA suggests early coupling of transcription-translation. Nat Commun 16, 6906 (2025).

29. Marshall, R.A., Aitken, C.E. & Puglisi, J.D. GTP hydrolysis by IF2 guides progression of the ribosome into elongation. Mol Cell 35, 37–47 (2009).

30. Ling, C. & Ermolenko, D.N. Initiation factor 2 stabilizes the ribosome in a semirotated conformation. Proc Natl Acad Sci U S A 112, 15874–9 (2015).

31. Myasnikov, A.G. et al. Conformational transition of initiation factor 2 from the GTP- to GDP-bound state visualized on the ribosome. Nat Struct Mol Biol 12, 1145–9 (2005).

32. Loveland, A.B., Demo, G. & Korostelev, A.A. Cryo-EM of elongating ribosome with EF-Tu*GTP elucidates tRNA proofreading. Nature 584, 640–645 (2020).

33. Carbone, C.E. et al. Time-resolved cryo-EM visualizes ribosomal translocation with EF-G and GTP. Nat Commun 12, 7236 (2021).

34. Pavlov, M.Y., Antoun, A., Lovmar, M. & Ehrenberg, M. Complementary roles of initiation factor 1 and ribosome recycling factor in 70S ribosome splitting. EMBO J 27, 1706–17 (2008).

35. Roske, J.J. et al. Structure of the 30S translation initiation complex coupled to paused RNA polymerase and its potential for riboregulation. Nat Commun 17, 693 (2025).

36. Pasqualato, S. & Cherfils, J. Crystallographic evidence for substrate-assisted GTP hydrolysis by a small GTP binding protein. Structure 13, 533–40 (2005).

37. Molt, R.W., Jr., Pellegrini, E. & Jin, Y. A GAP-GTPase-GDP-P(i) Intermediate Crystal Structure Analyzed by DFT Shows GTP Hydrolysis Involves Serial Proton Transfers. Chemistry 25, 8484–8488 (2019).

38. Pulk, A., Maivali, U. & Remme, J. Identification of nucleotides in E. coli 16S rRNA essential for ribosome subunit association. RNA 12, 790–6 (2006).

39. Hennelly, S.P. et al. A time-resolved investigation of ribosomal subunit association. J Mol Biol 346, 1243–58 (2005).

40. Parajuli, N.P., Emmerich, A., Mandava, C.S., Pavlov, M.Y. & Sanyal, S. Antibiotic thermorubin tethers ribosomal subunits and impedes A-site interactions to perturb protein synthesis in bacteria. Nat Commun 14, 918 (2023).

41. James, N.R., Brown, A., Gordiyenko, Y. & Ramakrishnan, V. Translational termination without a stop codon. Science 354, 1437–1440 (2016).

42. Qin, D. & Fredrick, K. Control of translation initiation involves a factor-induced rearrangement of helix 44 of 16S ribosomal RNA. Mol Microbiol 71, 1239–49 (2009).

43. Qin, D., Liu, Q., Devaraj, A. & Fredrick, K. Role of helix 44 of 16S rRNA in the fidelity of translation initiation. RNA 18, 485–95 (2012).

44. Carter, A.P. et al. Crystal structure of an initiation factor bound to the 30S ribosomal subunit. Science 291, 498–501 (2001).

45. Liu, Q. & Fredrick, K. Roles of helix H69 of 23S rRNA in translation initiation. Proc Natl Acad Sci U S A 112, 11559–64 (2015).

46. Ali, I.K., Lancaster, L., Feinberg, J., Joseph, S. & Noller, H.F. Deletion of a conserved, central ribosomal intersubunit RNA bridge. Mol Cell 23, 865–74 (2006).

47. Huang, C., Mandava, C.S. & Sanyal, S. The ribosomal stalk plays a key role in IF2-mediated association of the ribosomal subunits. J Mol Biol 399, 145–53 (2010).

48. Caban, K., Pavlov, M., Ehrenberg, M. & Gonzalez, R.L., Jr. A conformational switch in initiation factor 2 controls the fidelity of translation initiation in bacteria. Nat Commun 8, 1475 (2017).

49. Ge, X., Mandava, C.S., Lind, C., Aqvist, J. & Sanyal, S. Complementary charge-based interaction between the ribosomal-stalk protein L7/12 and IF2 is the key to rapid subunit association. Proc Natl Acad Sci U S A 115, 4649–4654 (2018).

50. Castro-Roa, D. & Zenkin, N. In vitro experimental system for analysis of transcription-translation coupling. Nucleic Acids Res 40, e45 (2012).

51. Mandava, C.S. et al. Bacterial ribosome requires multiple L12 dimers for efficient initiation and elongation of protein synthesis involving IF2 and EF-G. Nucleic Acids Res 40, 2054–64 (2012).

52. Svidritskiy, E. & Korostelev, A.A. Conformational Control of Translation Termination on the 70S Ribosome. Structure 26, 821–828 e3 (2018).

53. Fei, J. et al. A highly purified, fluorescently labeled in vitro translation system for single-molecule studies of protein synthesis. Methods Enzymol 472, 221–59 (2010).

54. Yokogawa, T., Kitamura, Y., Nakamura, D., Ohno, S. & Nishikawa, K. Optimization of the hybridization-based method for purification of thermostable tRNAs in the presence of tetraalkylammonium salts. Nucleic Acids Res 38, e89 (2010).

55. Gamper, H. & Hou, Y.M. A Label-Free Assay for Aminoacylation of tRNA. Genes (Basel) 11(2020).

56. Wang, W. et al. Loss of a single methylation in 23S rRNA delays 50S assembly at multiple late stages and impairs translation initiation and elongation. Proc Natl Acad Sci U S A 117, 15609–15619 (2020).

57. Punjani, A., Rubinstein, J.L., Fleet, D.J. & Brubaker, M.A. cryoSPARC: algorithms for rapid unsupervised cryo-EM structure determination. Nat Methods 14, 290–296 (2017).

58. Heymann, J.B. & Belnap, D.M. Bsoft: image processing and molecular modeling for electron microscopy. J Struct Biol 157, 3–18 (2007).

59. Mastronarde, D.N. Automated electron microscope tomography using robust prediction of specimen movements. J Struct Biol 152, 36–51 (2005).

60. Zheng, S.Q. et al. MotionCor2: anisotropic correction of beam-induced motion for improved cryo-electron microscopy. Nat Methods 14, 331–332 (2017).

61. Grant, T., Rohou, A. & Grigorieff, N. cisTEM, user-friendly software for single-particle image processing. Elife 7(2018).

62. Rohou, A. & Grigorieff, N. CTFFIND4: Fast and accurate defocus estimation from electron micrographs. J Struct Biol 192, 216–21 (2015).

63. Grigorieff, N. Frealign: An Exploratory Tool for Single-Particle Cryo-EM. Methods Enzymol 579, 191–226 (2016).

64. Ludtke, S.J. Single-Particle Refinement and Variability Analysis in EMAN2.1. Methods Enzymol 579, 159–89 (2016).

65. Pettersen, E.F. et al. UCSF Chimera--a visualization system for exploratory research and analysis. J Comput Chem 25, 1605–12 (2004).

66. Rybak, M.Y. & Gagnon, M.G. Structures of the ribosome bound to EF-Tu-isoleucine tRNA elucidate the mechanism of AUG avoidance. Nat Struct Mol Biol 31, 810–816 (2024).

67. Jumper, J. et al. Highly accurate protein structure prediction with AlphaFold. Nature 596, 583–589 (2021).

68. Emsley, P. & Cowtan, K. Coot: model-building tools for molecular graphics. Acta Crystallogr D Biol Crystallogr 60, 2126–32 (2004).

69. Liebschner, D. et al. Macromolecular structure determination using X-rays, neutrons and electrons: recent developments in Phenix. Acta Crystallogr D Struct Biol 75, 861–877 (2019).

70. Williams, C.J. et al. MolProbity: More and better reference data for improved all-atom structure validation. Protein Sci 27, 293–315 (2018).

71. Afonine, P.V. et al. New tools for the analysis and validation of cryo-EM maps and atomic models. Acta Crystallogr D Struct Biol 74, 814–840 (2018).

