## SUPPLEMENTARY DATA for "Molecular mechanism of IF1- and IF2-driven translation initiation in bacteria"

**The Supplementary data include:**

Supplementary tables 1-3

Supplementary figures 1-14

**Supplementary Table 1. Refinement statistics for cryo-EM structures 30S PIC-A, 30S PIC-B, 70S IC*, 70S IC-I, 70S IC-II and 70S EC.**

| | 30S PIC-A | 30S PIC-B | 70S IC* | 70S IC-I | 70S IC-II | 70S EC |
|---|---|---|---|---|---|---|
| **PDBID** | 9TKN | 9TKO | 9ZN2 | 9TKP | 9TKQ | 9TKR |
| **EMDB** | EMD-56034 | EMD-56035 | EMD-74431 | EMD-56036 | EMD-56037 | EMD-56038 |
| **Data collection and processing** | | | | | | |
| Magnification | 105,000x | 105,000x | 105,000x | 105,000x | 105,000x | 105,000x |
| Voltage (kV) | 300 | 300 | 300 | 300 | 300 | 300 |
| Electron exposure ($e^{-}/Å^2$) | 40 | 40 | 40.3 | 40 | 40 | 40 |
| Defocus range (μm) | -0.5 to -2.1 | -0.5 to -2.1 | -0.7 to -2.0 | -0.5 to -2.1 | -0.5 to -2.1 | -0.5 to -2.1 |
| Pixel size (Å) | 0.834 | 0.834 | 0.862 | 0.834 | 0.834 | 0.834 |
| Symmetry imposed | C1 | C1 | C1 | C1 | C1 | C1 |
| Initial particle (no.) | 351,980 | 351,980 | 757,723 | 825,112 | 825,112 | 825,112 |
| Final particle (no.) | 22,197 | 28,862 | 64,469 | 12,354 | 8,740 | 31,752 |
| Map resolution (Å) | 3.0 | 3.0 | 3.0 | 3.1 | 3.1 | 2.9 |
| FSC threshold | 0.143 | 0.143 | 0.143 | 0.143 | 0.143 | 0.143 |
| **Refinement** | | | | | | |
| Initial model used (PDB code) | 6WDD | 6WDD | 8G7P | 6WDD | 6WDD | 6WDD |
| Model resolution (Å) | 3.2 | 3.2 | 3.1 | 3.2 | 3.2 | 3.2 |
| Correlation Coefficient (cc_mask)* | 0.90 | 0.90 | 0.89 | 0.88 | 0.85 | 0.89 |
| Map sharpening *B* factor ($Å^2$) | -20 | -20 | -52.6 | -40 | -20 | -40 |
| Model composition* | | | | | | |
| Non-hydrogen atoms | 58,742 | 58,190 | 145,469 | 150,227 | 149,294 | 145,728 |
| Protein residues | 3,061 | 2,992 | 6,330 | 6,489 | 6,364 | 5,893 |
| RNA residues | 1,621 | 1,621 | 4,566 | 4,644 | 4,646 | 4,646 |
| *B* factors ($Å^2$)* | | | | | | |
| Protein | 143.19 | 145.83 | 53.09 | 150.19 | 147.73 | 131.25 |
| RNA | 131.76 | 132.18 | 60.30 | 134.97 | 134.63 | 121.78 |
| GTP/GDP/GDPCP | 293,12 | 264.96 | 30.30 | 135.11 | 157.64 | - |
| $Mg^{2+}$ | 292.86 | 276.61 | 39.27 | 156.23 | - | - |
| R.m.s. deviations*,§ | | | | | | |
| Bond lengths (Å) | 0.005 | 0.004 | 0.008 | 0.005 | 0.005 | 0.004 |
| Bond angles (°) | 0.78 | 0.76 | 0.70 | 0.75 | 0.78 | 0.73 |
| Validation# | | | | | | |
| MolProbity score | 1.82 | 1.82 | 1.37 | 1.5 | 1.87 | 1.71 |
| Clashscore | 4.98 | 5.34 | 2.39 | 1.96 | 5.59 | 4.12 |
| Poor rotamers (%) | 0.2 | 0.1 | 0.0 | 0.0 | 0.1 | 0.0 |
| Ramachandran plot# | | | | | | |
| Favored (%) | 89.53 | 90.58 | 94.92 | 90.4 | 89.18 | 91.09 |
| Allowed (%) | 10.27 | 9.28 | 5.00 | 9.51 | 10.61 | 8.76 |
| Disallowed (%) | 0.20 | 0.14 | 0.08 | 0.09 | 0.21 | 0.12 |
| Validation (RNA)# | | | | | | |
| Good sugar pucker (%) | 99.1 | 99.3 | 99.7 | 99.3 | 99.2 | 99.3 |
| Good backbone (%) | 84.9 | 84.7 | 84.6 | 83.9 | 82.3 | 84.2 |

*from Phenix

### from Molprobity

§ root mean square deviations

**Supplementary Table 2. Comparison of buried surface area (BSA) in 70S IC-I and 70S IC***

| Bridges | $BSA_{70S\ IC\text{-}I}$ (Å$^2$) | $BSA_{70S\ IC*}$ (Å$^2$) | BSA loss (Å$^2$) | % BSA loss | % BSA retained |
|---|---|---|---|---|---|
| B1a[1] | 166.3 | 61.9 | 104.4 | 62.8 | 37.2 |
| B1b[1] | 203.6 | 121.9 | 81.7 | 40.1 | 59.8 |
| **B2a** | 1094.3 | 15.3 | 1079.0 | 98.6 | 1.4 |
| **B2b** | 190.2 | 26.0 | 164.2 | 86.3 | 13.7 |
| B3 | 437.6 | 357.5 | 80.1 | 18.3 | 81.6 |
| B4 | 408.9 | 430.8 | - | - | 105.3 |
| B5 | 639.6 | 670.2 | - | - | 104.8 |
| B6 | 466.9 | 477.2 | - | - | 102.2 |
| **B7a** | 324.8 | 130.5 | 194.3 | 59.8 | 40.2 |
| **B7b** | 242.4 | 76.8 | 165.6 | 68.3 | 31.7 |
| B8 | 482.8 | 476.1 | 6.7 | 14 | 98.6 |

[1] Bridges B1a and B1b involve regions in the head domain of the 30S subunit, which are often disordered and therefore, the BSA calculations are considered unreliable.

**Supplementary Table 3. Buffer compositions used for purification of IFs and IF2 variants for kinetic and cryo-EM studies.**

| **Kinetic studies** | |
|---|---|
| *Buffer* | *Composition* |
| Buffer A | 20 mM Tris-HCl pH 7.5, 300 mM NaCl, 5 mM $MgCl_2$, 10% glycerol |
| Buffer B | Buffer A + 300 mM imidazole |
| **Cryo-EM studies** | |
| *Buffer* | *Composition* |
| Buffer A - IF1 | 10 mM Tris-HCl pH 7.5, 50 mM NaCl, 10 mM $MgCl_2$, 30 mM imidazole, 5% (v/v) glycerol, 6 mM βME + EDTA-free protease inhibitors (Complete Mini, Merck) |
| Buffer A - IF2α | 20 mM Tris-HCl pH 8.0, 200 mM NaCl, 5 mM $MgCl_2$, 3mM imidazole, 6 mM βME + EDTA-free protease inhibitors (Complete Mini, Merck) |
| Buffer B - IF1 | Buffer A + 500 mM imidazole |
| Buffer B - IF2α | Buffer A + 200 mM imidazole |
| Buffer C - IF1 | 10 mM Tris-HCl pH 7.5, 50 mM NaCl, 10 mM $MgCl_2$, 6 mM βME + EDTA-free protease inhibitors (Complete Mini, Merck) |
| Buffer C - IF2α | 40 mM Tris-HCl pH 7.5, 30 mM NaCl, 40 mM $NH_4Cl$, 5 mM $MgCl_2$, 6 mM βME + EDTA-free protease inhibitors (Complete Mini, Merck) |
| Buffer E - IF1 | 10 mM Tris-HCl pH 7.5, 80 mM NaCl, and 10 mM $MgCl_2$ |
| Buffer E - IF2α | 40 mM Tris-HCl pH 7.5, 80 mM NaCl, 40 mM $NH_4Cl$, 5 mM $MgCl_2$ |
| Storage buffer - IF1 | 10 mM Tris-HCl pH 7.5, 80 mM NaCl, 10 mM $MgCl_2$, 10% (v/v) glycerol |
| Storage buffer - IF2α | 40 mM Tris-HCl pH 7.5, 80 mM NaCl, 40 mM $NH_4Cl$, 5 mM $MgCl_2$, 0.5 mM EDTA and 15% (v/v) glycerol |

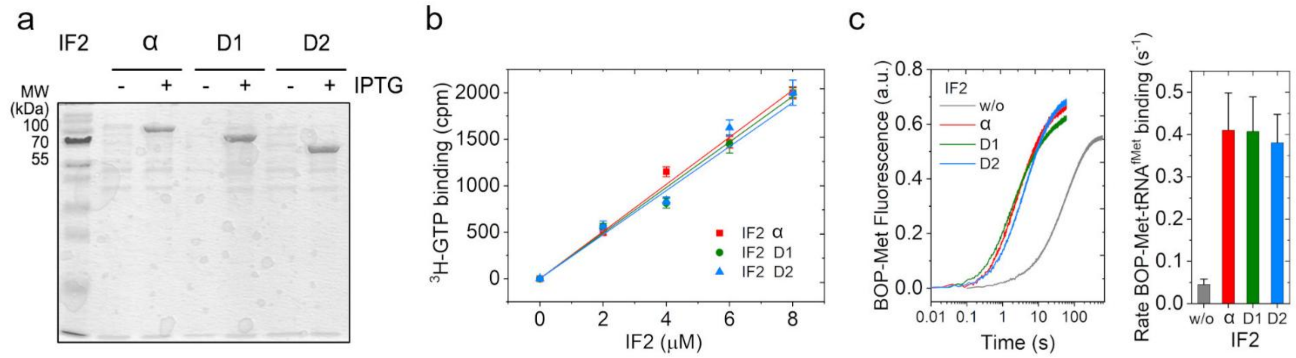


**Supplementary Figure 1. Purification and biochemical characterization of the IF2 variants.** (**a**) SDS-PAGE of IPTG induced overexpression of IF2α and N-terminal truncation mutants D1 (Δ1–159) and D2 (Δ1–289), respectively. The over-expressed protein bands correspond to the respective sizes. (**b**) [$^3$H]-GTP binding assay for IF2 variants. IF2 variants (IF2α, D1 or D2) (0 – 8 μM) were mixed with 100 μM [$^3$H]-GTP and incubated at 37 °C for 10 minutes to allow binding. The reactions were quenched with 50% ice-cold formic acid and filtered through nitrocellulose filter. The filters were transferred to Beckman® Liquid Scintillation Counter and the radioactive counts were measured. The cpm for [$^3$H]-GTP/GDP was plotted against respective IF2 concentration. The plots showed similar linear increase of [$^3$H]-GTP/GDP counts for increase in IF2 concentration. The IF2α and the D1 and D2 mutants exhibit comparable GTP binding, indicating that truncation of the N-terminal region does not influence the GTP binding activity of the factor. (**c**) Kinetics of BOP•Met-tRNA$^{fMet}$ binding to the ribosome in the presence of IF2 variants. The binding was followed in stopped-flow at 590 nm ($\lambda_{Ex}$ = 560 nm). Similar BOP•Met-tRNA$^{fMet}$ binding kinetics with all IF2 variants suggest that N-terminal truncation of IF2 does not affect initiator tRNA recruitment. All kinetic data are fitted with OriginPro 2016 (Origin LabCorp) and rates ± s.d. are estimated from three independent experiments.

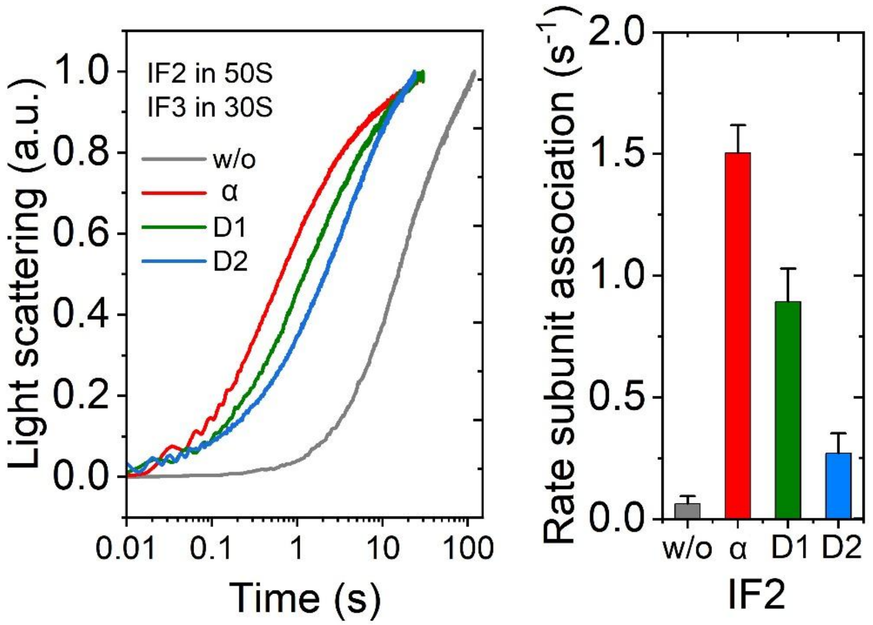


**Supplementary Figure 2. Comparison of IF2α, D1 and D2 variants in subunit association in the presence of IF3.** Kinetics of association of 50S subunit and 30S PIC containing IF1 and IF3, followed by Rayleigh light scattering in stopped-flow (left panel). The IF2 variants were supplied with 50S subunit so that the reactions will be rate-limited by and would reflect binding of the IF2 variants to the 30S PIC. The right panels demonstrate the rates of subunit association for each IF2 construct. Data are presented as mean ± s.d. from three independent experiments. The rates summarized in the right panel present that they show a pattern of IF2α > D1 > D2 similar to the reaction without IF3 although the overall rates decreased significantly. The conclusion, thus, is that the binding of the IF2 variants and the subsequent subunit association are not affected by the presence of IF3.

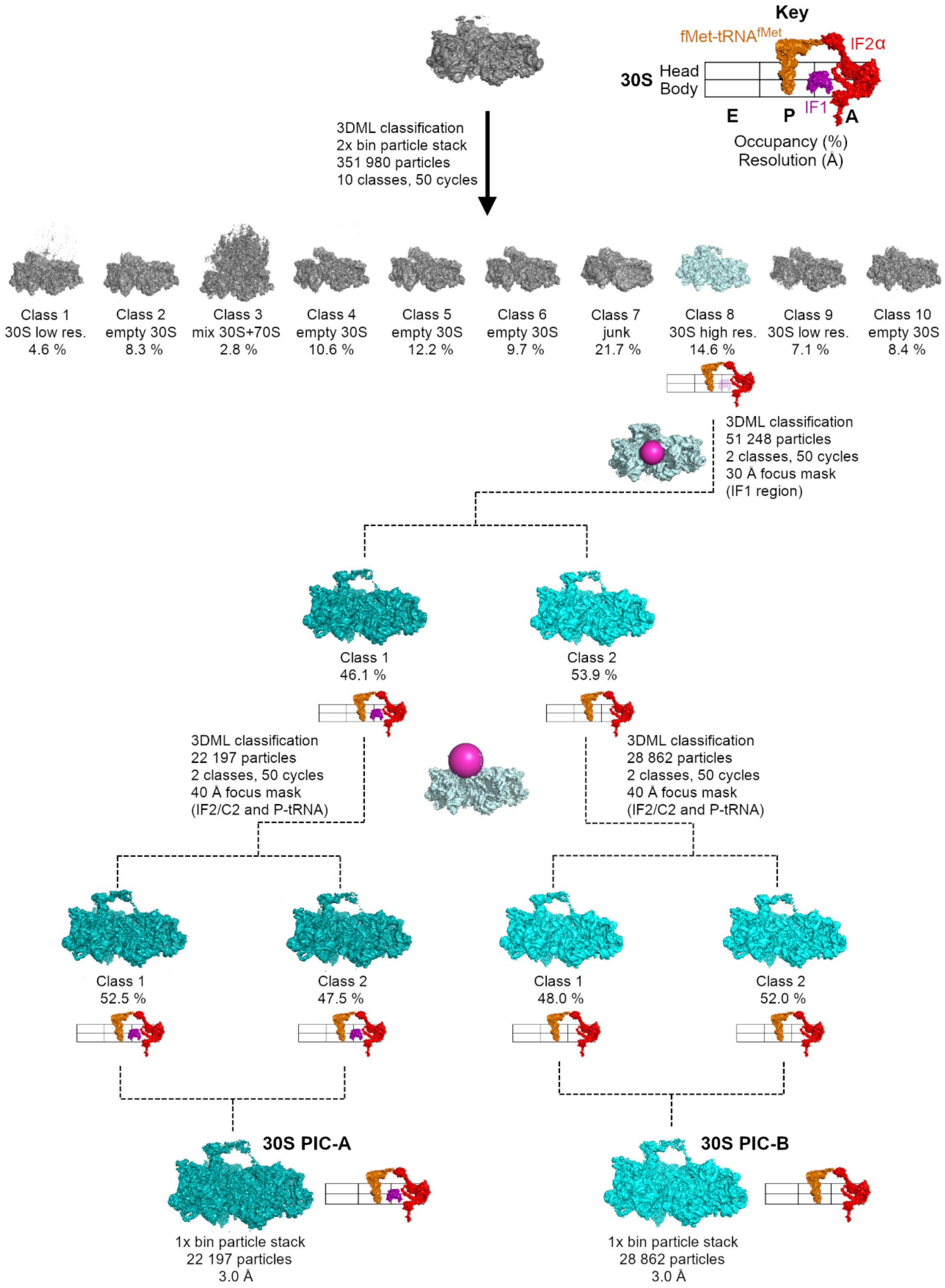


**Supplementary Figure 3. Ensemble cryo-EM data processing and classification scheme for the 30S PIC structures.** Low-resolution and junk classes are shown in grey, while classes advancing to 3D classification are shown in dark cyan (30S PIC-A) and light cyan (30S PIC-B). Final maps used for structural modelling are displayed at the bottom, along with the number of particles and corresponding resolutions (FSC = 0.143).

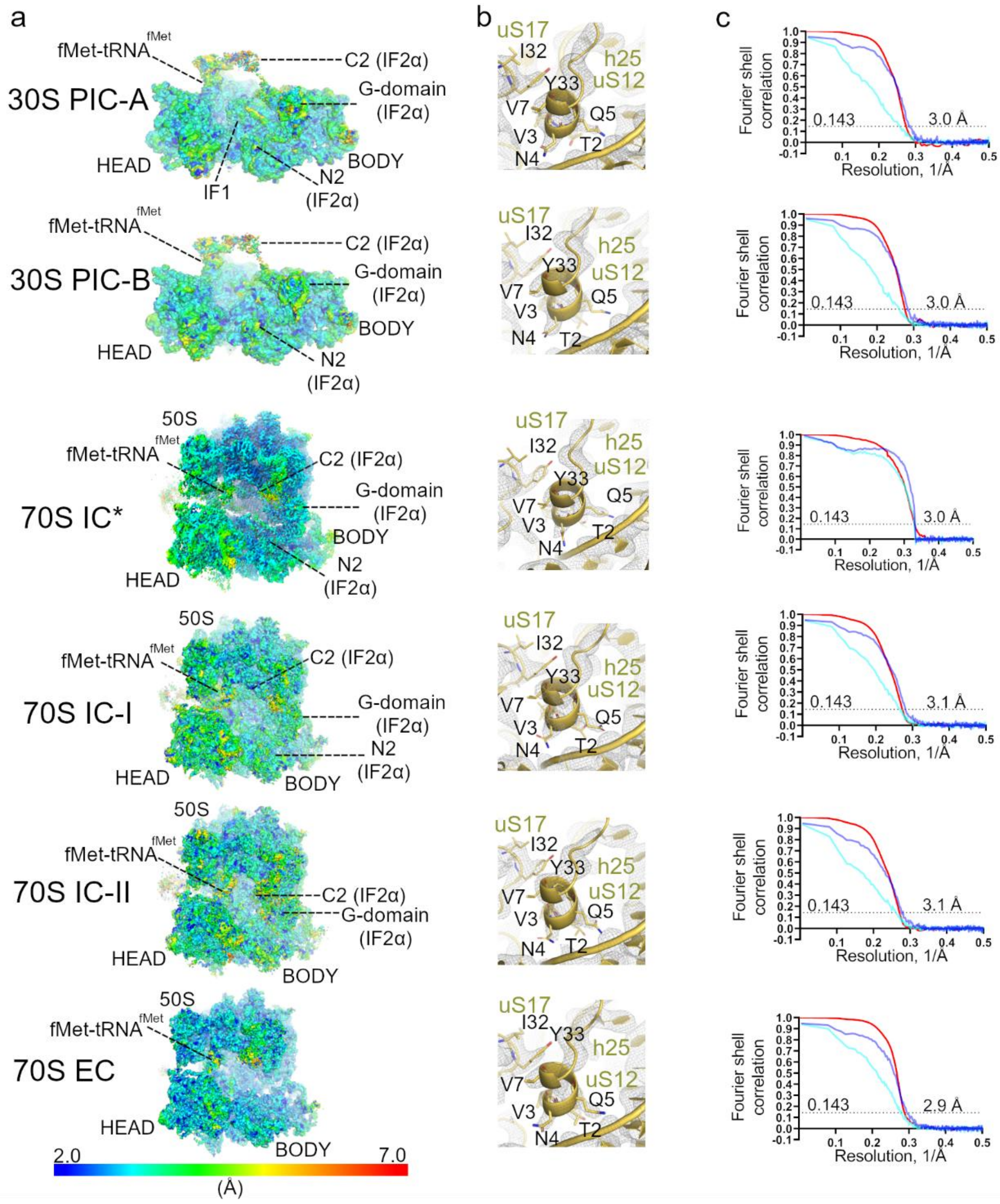


**Supplementary Figure 4. Global and local resolution for 30S PIC-A, 30S PIC-B, 70S IC*, 70S IC-I, 70S IC-II and 70S EC.** (**a**) Local resolutions in the cryo-EM maps are shown using a colour gradient ranging from 2.0 to 7.0 Å. (**b**) Example of local map resolution for 30S PIC-A, 30S PIC-B, 70S IC*, 70S IC-I, 70S IC-II and 70S EC in the region of the uS12 of the 30S subunit. The selected residues of uS12 are shown in stick representation. The original non-sharpened maps (gray mesh) are shown at 6.0 σ. (**c**) Fourier shell correlation (FSC) between even- and odd-particle half maps (red) show that map resolutions range from 2.9 to 3.1 Å (at FSC = 0.143, dotted line); FSC between final models and final maps (cyan) masked and unmasked FSCs are also shown.

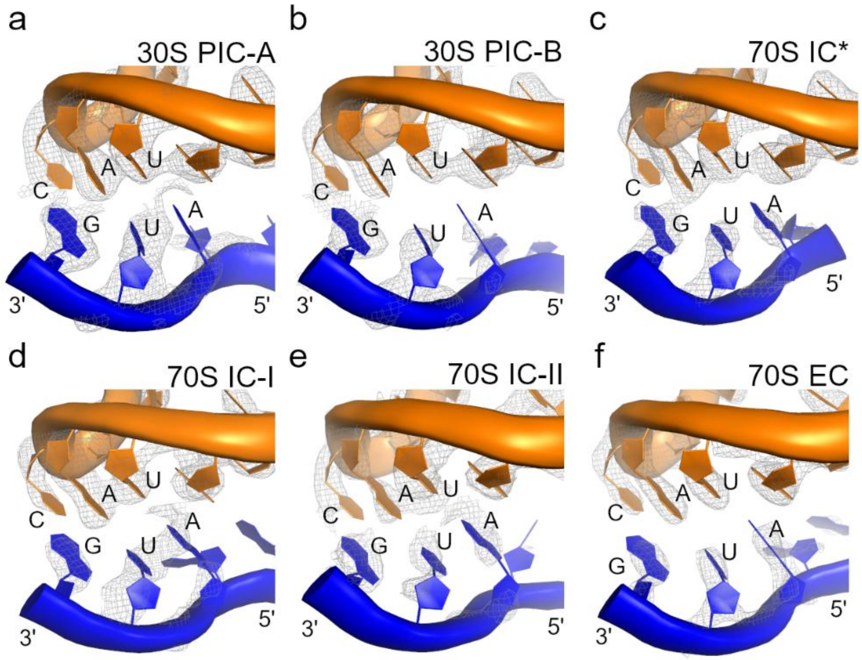


**Supplementary Figure 5. Base pairing between the mRNA start codon and the initiator tRNA anticodon.** Close-up views of the mRNA AUG start codon (shown in blue) and the anticodon of fMet-tRNA$^{fMet}$ (shown in orange) positioned within the P-site of the 30S subunit. The original cryo-EM density maps are displayed as grey mesh and were segmented to enhance visualization of both the tRNA and mRNA regions. (**a**) For 30S PIC-A, the cryo-EM map was contoured at 4.0 σ for fMet-tRNA$^{fMet}$ and 2.4 σ for mRNA. (**b**) For 30S PIC-B, the cryo-EM map was contoured at 4.2 σ for fMet-tRNA$^{fMet}$ and 2.4 σ for mRNA. (**c**) For 70S IC*, the cryo-EM map was contoured at 2.8 σ for fMet-tRNA$^{fMet}$ and mRNA. (**d**) For 70S IC-I, the cryo-EM map was contoured at 3.5 σ for fMet-tRNA$^{fMet}$ and 2.3 σ for mRNA. (**e**) For 70S IC-II, the cryo-EM map was contoured at 3.0 σ for fMet-tRNA$^{fMet}$ and 1.8 σ for mRNA. (**f**) For 70S EC, the cryo-EM map was contoured at 3.5 σ for fMet-tRNA$^{fMet}$ and 2.1 σ for mRNA.

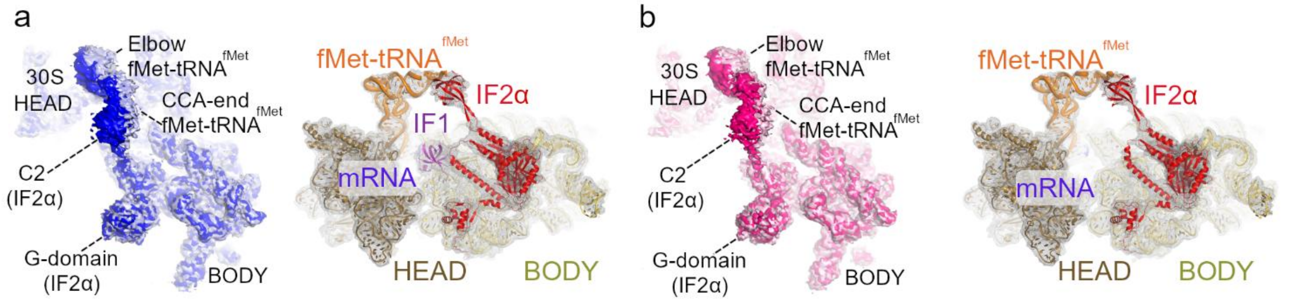


**Supplementary Figure 6. Conformational variability of the IF2α C2-domain and the 3′-CCA end of initiator tRNA in the PIC structures.** (**a**) 30S PIC-A: Left panel shows the superposition of two cryo-EM surfaces obtained from subclassification, highlighting conformational variability in the IF2α C2-domain and the 3′-CCA end of fMet-tRNA$^{fMet}$. As no additional structural differences were observed between the two subclasses, and no improvement in resolution was achieved in the C2-domain region, the subclasses were merged to generate the final 30S PIC-A reconstruction. Right panel shows the final 30S PIC-A model fitted into the corresponding low-pass filtered cryo-EM map (grey mesh; B-factor 50 Å$^2$ and contour level 3.5 σ), marking the position of the C2-domain and the 3′-CCA end of the initiator tRNA. (**b**) 30S PIC-B: as in panel (a), the left panel shows two superimposed surfaces from subclassification, revealing conformational variability in the same regions. As with 30S PIC-A, the lack of additional differences and insufficient resolution gain led to the merging of the subclasses to form the final 30S PIC-B structure. The right panel shows the 30S PIC-B model fitted into its corresponding low-pass filtered cryo-EM map (grey mesh; B-factor 50 Å$^2$ and contour level 3.5 σ), marking the position of the C2-domain and the 3′-CCA end of the initiator tRNA.

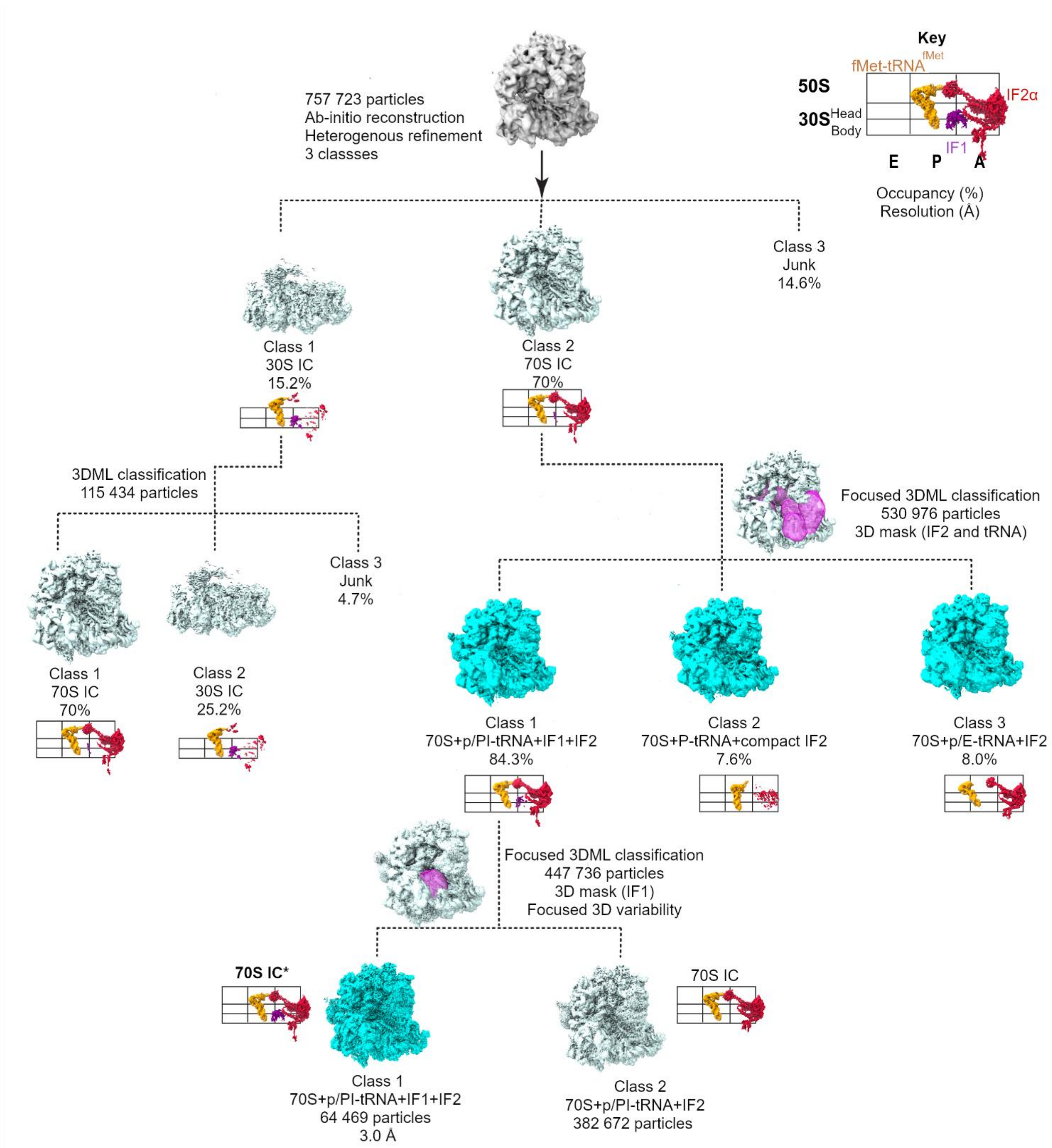


**Supplementary Figure 7. Cryo-EM data processing and classification scheme for 70S IC*.** Classes that advanced to further 3D classification are coloured in dark cyan. The final map used for structural modelling is shown at the bottom in dark cyan, along with the corresponding particle number and final resolution (FSC = 0.143).

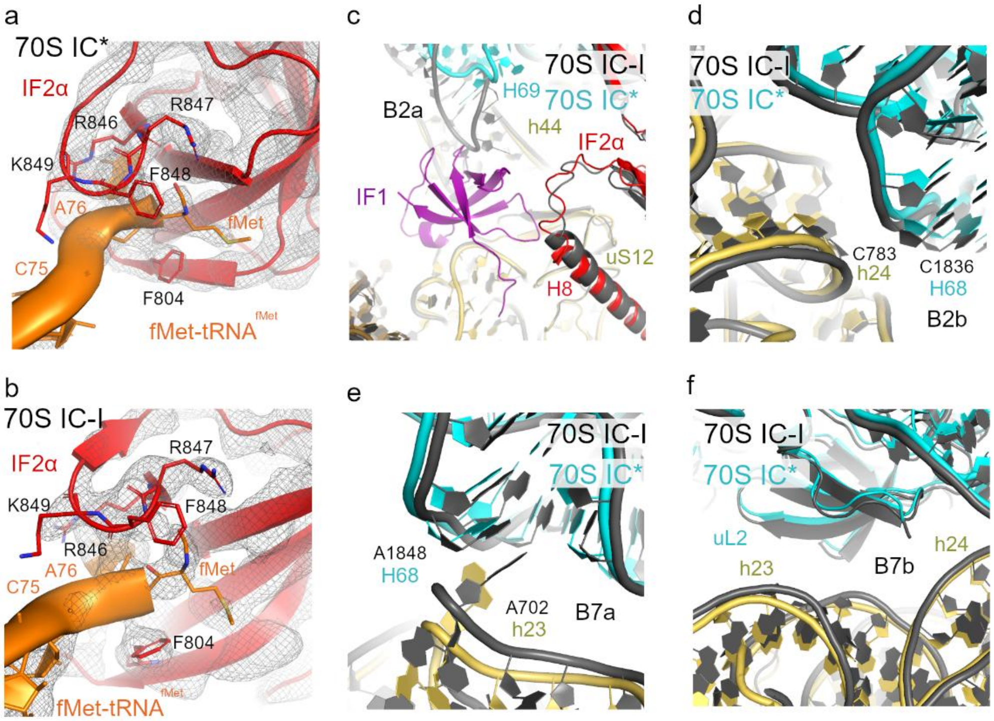


**Supplementary Figure 8. Stabilization of initiator tRNA by IF2α and formation of inter-subunit bridges.** Interface between the IF2α C2-domain and the 3′-CCA end of fMet-tRNA$^{fMet}$ in 70S IC* (**a**) and 70S IC-I (**b**). The cryo-EM density is shown as a grey mesh contoured at 2.2 σ in (a) and at 2.0 σ in (b). The fMet moiety and nearby C2-domain residues are shown as sticks. Close-up structural comparisons of 70S IC* and 70S IC-I highlighting incompletely formed inter-subunit bridges B2a (**c**), B2b (**d**), B7a (**e**), and B7b (**f**). Structures were aligned on the 16S rRNA of the 30S body, with 70S IC-I shown in dark grey.

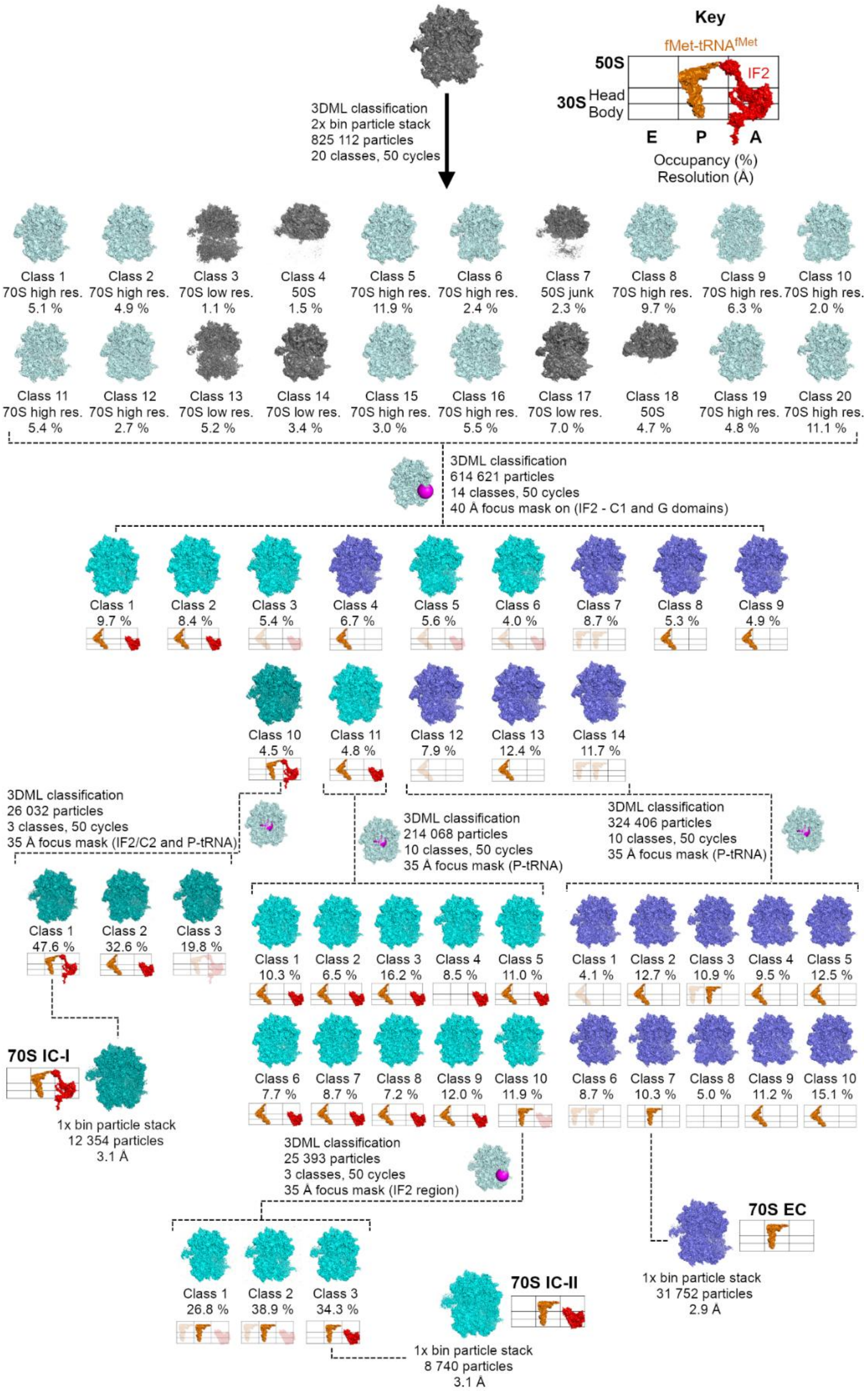


**Supplementary Figure 9. Ensemble cryo-EM data processing and classification scheme for 70S IC and 70S EC particles.** Junk and low-resolution classes are shown in grey, while classes that advanced to further 3D classification are coloured dark cyan (70S IC-I), light cyan (70S IC-II), and navy blue (70S EC). Final maps used for structural modelling are shown at the bottom, along with the corresponding particle numbers and final resolutions (FSC = 0.143).

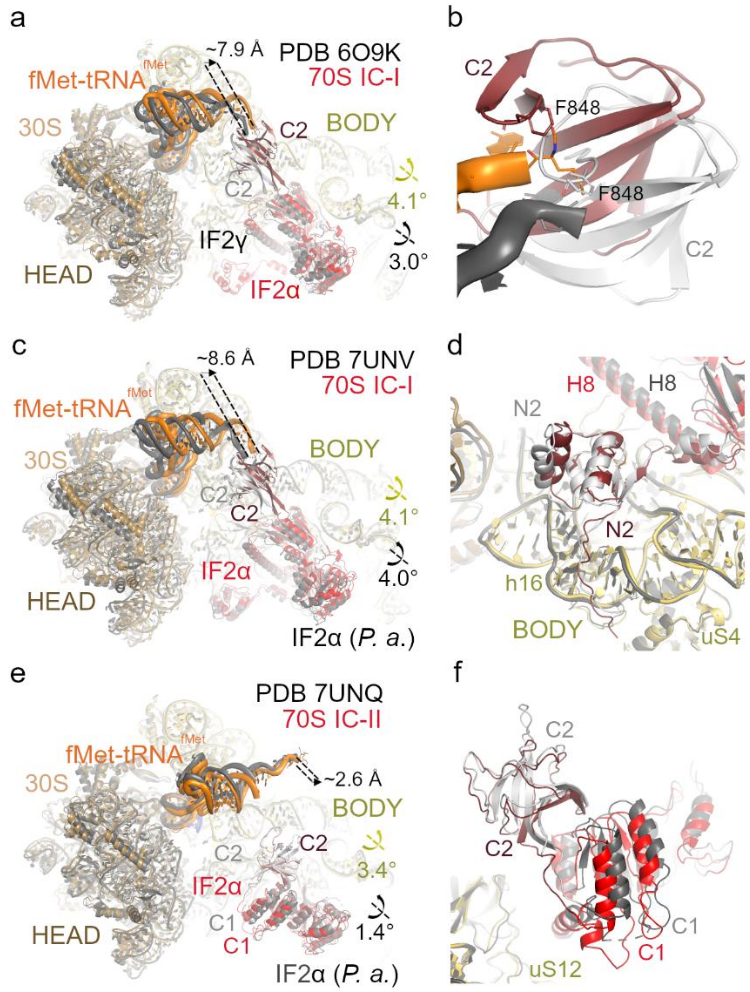


**Supplementary Figure 10. Structural comparison of 70S IC-I and 70S IC-II with previously reported cryo-EM structures of the 70S initiation complex.** Structural alignments were performed based on the 30S body 16S rRNA for each pairwise comparison. (**a**) Superposition of 70S IC-I and *E. coli* 70S IC (PDB 6O9K), showing 30S body rotation. Rotation angles are indicated by yellow (70S IC-I) and black (PDB 6O9K) arrows. The positional variability between the CCA-ends of fMet-tRNA$^{fMet}$ (orange - 70S IC-I, dark grey - 6O9K) is shown as a difference in distance indicated by arrow. (**b**) Close-up view of the IF2`s C2-domain and fMet-tRNA$^{fMet}$ interface between 70S IC-I and PDB 6O9K, highlighting the positional displacement between Phe848 residue located in C2-domain. (**c**) Superposition of 70S IC-I and *P. aeruginosa* structure II-A (PDB 7UNV), showing 30S body rotation. Rotation angles are indicated by yellow (70S IC-I) and black (PDB 7UNV) arrows. The positional variability between the CCA-ends of fMet-tRNA$^{fMet}$ (orange - 70S IC-I, dark grey - 7UNV) is shown as a difference in distance indicated by arrow. (**d**) Structural alignment of the IF2α N2-domain between 70S IC-I (ruby) and PDB 7UNV (light grey), both docked onto h16 of 16S rRNA. (**e**) Superposition of 70S IC-II and *P. aeruginosa* structure I-A (PDB 7UNQ), showing 30S body rotation. Rotation angles are indicated by yellow (70S IC-I) and black (PDB 7UNQ) arrows. (**f**) Close-up view of the compact conformation of IF2α in both 70S IC-II and PDB 7UNQ. In all panels, the 70S IC-I and IC-II structures are coloured as in Fig. 2, while the previously reported 70S initiation complexes are shown in shades of grey.

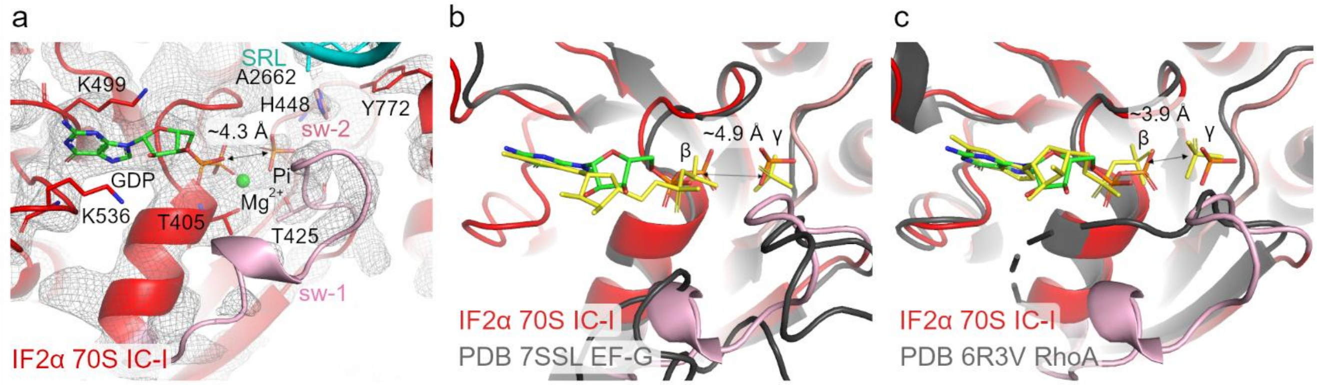


**Supplementary Figure 11. Structural comparison of the IF2α G-domain in the 70S IC-I and related GTPases.** (**a**) Close-up view of the IF2α G-domain active site in the 70S IC-I structure. The nucleotide-binding pocket is shown with the modelled GDP–Pi state, with coordinating residues displayed as sticks and the $Mg^{2+}$ ion shown as a green sphere. The catalytic His448 within the switch II (sw-2) region is highlighted, together with the switch I (sw-1) region and the sarcin–ricin loop (SRL) of the 23S rRNA. The cryo-EM density is shown as a grey mesh, contoured at 2.2 σ for IF2α and 4.0 σ for the SRL. The distance between the phosphorus atoms of the β- (GDP) and γ-phosphates (Pi) is indicated by a double-headed arrow (4.3 Å). Structural alignment of the IF2α G-domain from the 70S IC-I with representative GDP–Pi states of other GTPases, including (**b**) the ribosome-bound EF-G translocation complex (PDB 7SSL) and (**c**) a Ras-like GTPase (PDB 6R3V), shown in dark grey. The alignment reveals an active-site geometry, with similar positioning of the β- (GDP) and γ-phosphates (Pi) of these GTPases (shown as yellow sticks), supporting a catalytically competent configuration. Distances between the corresponding phosphate groups are indicated by double-headed arrows (EF-G; 4.9 Å and RhoA; 3.9 Å). The alignment was performed using the G-domain of IF2α.

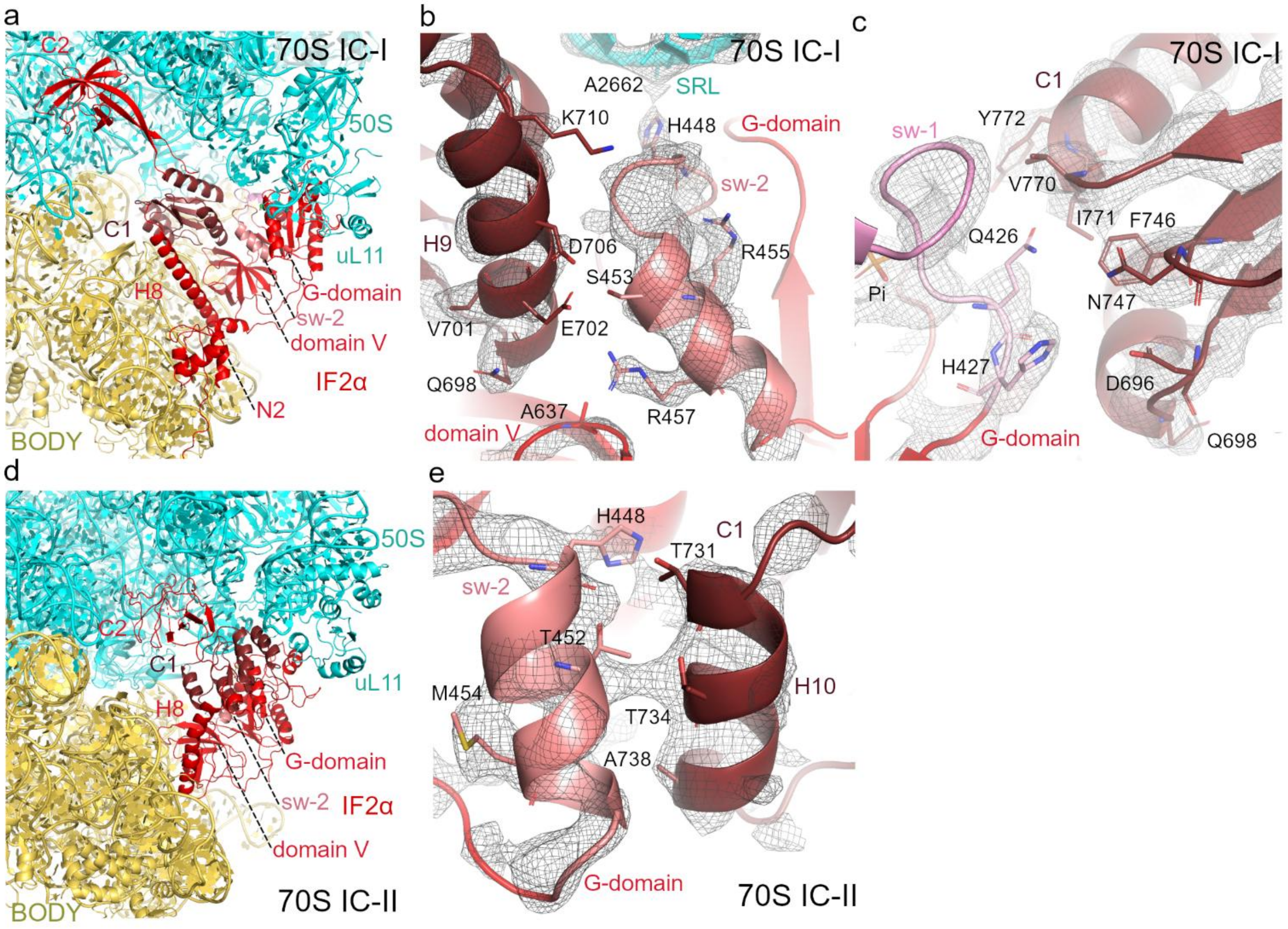


**Supplementary Figure 12. Relocation of the C2-domain and stabilization of the G-domain switch loops in 70S IC-I and 70S IC-II.** (**a**) Overall view on IF2α location in 70S IC-I, with highlighted IF2α domains respectively. (**b**) Interaction interface between IF2α sw-2 region in G-domain, and helix H9 of C1-domain and domain V. The Ser453 in sw-2 region is in close contact (~3.6 Å) with Glu702 located in H9 of C1-domain. The Arg457 in sw-2 is closely positioned (~3.6 Å) to Ala637 in domain V of IF2α. The original cryo-EM map is shown as a grey mesh and contoured at 2.0 σ. (**c**) Interaction interface between IF2α sw-1 region and C1-domain. The Gln426 in sw-1 is positioned next to the N-terminal extremity of helix H12 of C1-domain (Phe746, Val770 and Ile771). (**d**) Overall view on IF2α location in 70S IC-II, with highlighted IF2α domains and coloured as in panel (a). (**e**) Interaction interface between IF2α sw-2 and helix H10 of C1-domain. The Thr452 in sw-2 is in close contact (~3.1 Å) with Thr734 located in H10 of C1-domain. All panels use the same colour scheme as shown in Fig. 2.

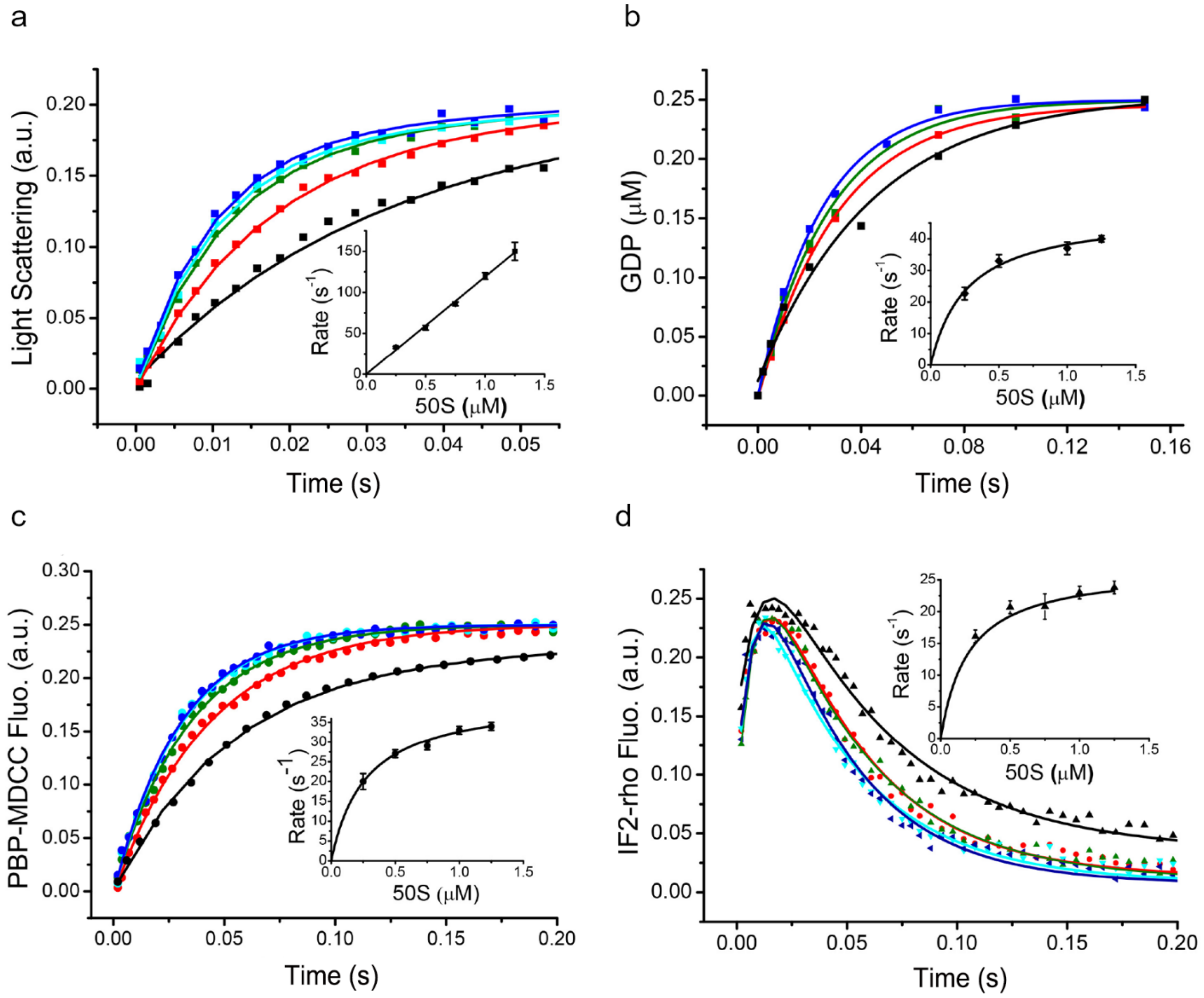


**Supplementary Figure 13. Fast kinetics measurements of translation initiation steps.** Kinetic analysis of subunit association (**a**), GTP hydrolysis (**b**), Pi release (**c**), and IF2α release (**d**) at 37ºC, measured at varying concentrations of the 50S subunit (0.25 μM, black trace; 0.5 μM, red trace; 0.75 μM, green trace; 1μM, cyan trace; and 1.25 μM, blue trace). The 30S pre-initiation complexes (30S PICs) were formed by incubating 30S, mRNA, fMet-tRNA$^{fMet}$, IF1, IF2α and GTP. The 30S PICs were rapidly mixed with 50S subunits using stopped-flow (a, c, and d) or quench-flow (b) techniques. For GTP hydrolysis (b), [$^3$H]-GTP was used instead of unlabeled GTP. For IF2α release (d), rhodamine-labeled IF2α (IF2-rho) was used in the 30S PICs in subunit association assays. The insets show Michaelis-Menten plots of the rate of each step as a function of 50S subunit concentration. Data are mean ± s.d. from three independent experiments.

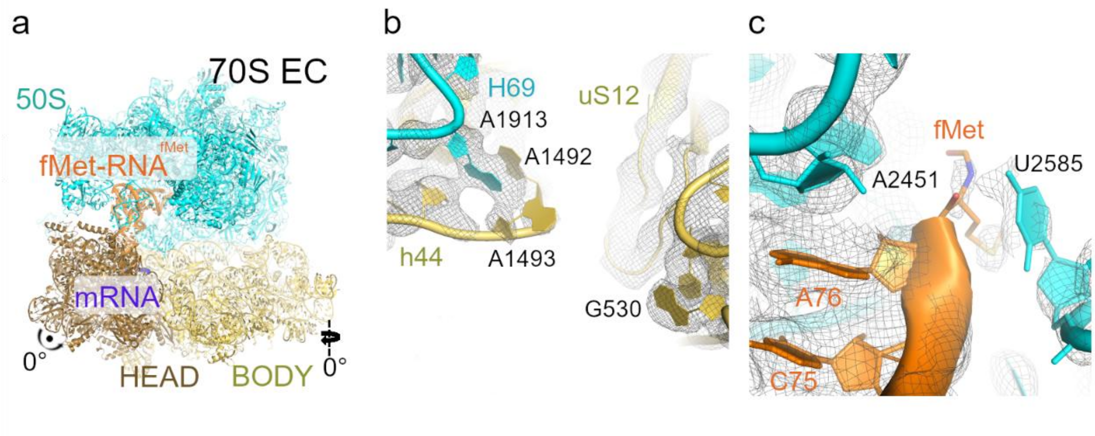


**Supplementary Figure 14. Cryo-EM structure of 70S EC.** (**a**) Overall structure view of the 70S elongation complex (EC). Arrows indicate the 30S head swivel and body rotation. (**b**) Close-up view of the DC in the 70S EC, showing the decoding nucleotides. The cryo-EM density is displayed as a grey mesh contoured at 2.5 σ. (**c**) Close-up view of the PTC, highlighting the 3′-CCA end of fMet-tRNA$^{fMet}$ in the P/P state, deeply buried within the PTC. The fMet moiety and surrounding 23S rRNA nucleotides are shown in stick representation. The cryo-EM map was segmented to display the fMet-tRNA$^{fMet}$ region at 1.4 σ and the 23S rRNA at 3.2 σ.
